# Sex differences in insular cortex function in persistent alcohol drinking despite aversion in mice

**DOI:** 10.1101/2023.10.04.560817

**Authors:** Claudia Fornari, Daria Ricci, Yoni Couderc, Carmen Guerrero-Márquez, Praneeth Namburi, Camille Penet, Céline Nicolas, Anna Beyeler

## Abstract

One major hallmark of alcohol use disorder (AUD) is the persistence of drinking despite negative consequences. Among indicators of AUD vulnerability, binge drinking has been identified as one of the strongest risk factors. Although the lifetime prevalence of both binge drinking and AUD has historically been higher in men than women, this gap has dramatically narrowed in the last decade. Additionally, sex differences in AUD and binge drinking have been found in clinical and preclinical studies. At the neurobiological level, the insular cortex plays an important role in AUD, with the anterior (aIC) and posterior (pIC) divisions supporting different functions. However, the contributions of the aIC and pIC sections in sexual dimorphism of alcohol binge drinking and the persistence of alcohol drinking despite aversion remain to be uncovered. Using the drinking in the dark model in mice, we validated that female mice have a higher binge ethanol intake compared to males. To evaluate persistent ethanol consumption despite aversion, we supplemented ethanol with the bitter compound quinine, and found a higher persistent drinking in females compared to males. Using fiber photometry recordings, we revealed that aIC activity was increased during binge and persistent ethanol consumption independently of sex, whereas pIC glutamatergic neuron activity was higher during persistent ethanol drinking, specifically in female mice. Using chemogenetics, we revealed that inhibition of aIC glutamatergic neurons reduced intake of bitter solutions independently of the solvent (ethanol or water) in both sexes. In addition, inhibition of pIC glutamatergic neurons exclusively reduced persistent ethanol drinking in females, while decreasing quinine consumption only in males. These findings suggest a sex-dependent function of the pIC in the persistence of ethanol consumption, providing a starting point in understanding sex-specific functions of the insular cortex in the neurobiology of AUD.

## INTRODUCTION

Alcohol use disorder (AUD) is a chronic relapsing disorder defined by the persistence of excessive alcohol consumption despite physical and psychological negative consequences (1), which is a considerable impediment to AUD treatment (2,3). AUD is a major public health burden, as 6% of the world population is affected by alcohol morbidity or mortality (4). Among the indicators of vulnerability to AUD, binge drinking has been identified as a strong risk factor (3,5–7). Binge drinking is a recreational pattern of alcohol use characterized by episodic and excessive consumption that raises blood alcohol concentration to 80 mg/dL or higher, typically within about two hours. A longitudinal study showed that 43% of adolescents (13 to 18 years old) who binge-drink alcohol developed AUD at the age of 21, whereas only 7% of non-bingers developed AUD at the same age (8). In preclinical studies, binge drinking is modeled using the drinking in the dark (DID) procedure, where mice have repeated access to alcohol for a limited time during the dark phase of the circadian rhythm (active period), which allows to reach blood ethanol concentration (BEC) of 80 mg/dL as observed in humans (9–11). One core symptom of drug addiction is the persistence of drug use despite negative consequences. This specific aspect can be modeled in animals with different approaches. Among them, the DID procedure can be adapted to study the persistence of alcohol drinking despite aversive outcomes by adulterating ethanol with quinine (e.g., 100-500µM), a bitter and aversive substance for mice (12–15).

Historically, men have been reported as more likely than women to drink alcohol and exhibit pathological drinking behaviors. In 2020, AUD lifetime prevalence was still lower in women than men, reaching respectively 23% and 36% (16). However, a recent longitudinal study showed that the increase rate of AUD over a decade was drastically higher in women than men, with respective rates of 84% and 35% (17). These epidemiological data highlight that the sex difference in AUD prevalence is narrowing. Additionally, women transition faster to AUD after regular or chronic alcohol consumption (18,19) and are subsequently more likely to develop alcohol-related diseases (e.g. cardiovascular and hepatic diseases, 19–21). Interestingly, preclinical studies revealed sex differences in alcohol intake as well (23), with female rodents drinking more than males across different models of intake, including binge drinking and drinking despite taste aversion (13,24–31). This higher propensity to alcohol drinking in females is proposed to be, at least in part, due to their lowest sensitivity to aversive properties of alcohol, as female rodents are more resistant to ethanol-conditioned taste aversion (32–34).

The insular cortex (or insula) is strongly involved in drug addiction (35–37), including AUD (38– 43). Indeed, human functional imaging studies in alcohol-dependent subjects have demonstrated that alcohol cues trigger greater activity responses in the insula compared to control neutral cues (44,45). Furthermore, insula white matter volume is correlated to binge drinking frequency in adolescents (46). Anatomically, the insula is subdivided into the anterior (aIC) and posterior (pIC) sections, which are proposed to have antagonistic functions in a wide range of behaviors (47–49). In AUD patients, structural studies report a reduction of the aIC volume and gray matter density compared to healthy controls, which is correlated with higher compulsive drinking measures (50– 53). In addition, aIC functional connectivity with several brain regions (e.g., hippocampus, medial orbitofrontal regions) is higher in patients with AUD compared to social drinkers (54) and healthy controls (55). Intriguingly, less information is known about the role of pIC in AUD. Indeed, only two recent studies reported a decrease in pIC gray matter volumes in AUD patients (55,56) and a higher resting state activity of the pIC relative to healthy controls (55), suggesting the pIC also plays a role in AUD. Preclinical studies confirmed the contribution of both aIC and pIC in ethanol-drinking behaviors. Indeed, chemogenetic activation or inhibition of aIC neurons respectively decreased and increased alcohol intake in several behavioral paradigms such as self-administration and intermittent 2-bottle choice (41,57,58). In contrast, pharmacological inactivation of pIC neurons decreased alcohol self-administration (59), suggesting divergent functions of aIC and pIC in alcohol drinking. However, how sex as a biological variable influences those divergent contributions remained untested.

Recent findings point towards structural sexual dimorphism of the insula, with a larger volume in men than women in physiological conditions (60), whereas in AUD long-term abstinent patients, this ratio is inverted in both right aIC and pIC (53). In support of these clinical findings, preclinical studies highlighted sex differences in insula functions in alcohol-drinking behaviors. For example, in alcohol binge-drinking mice, optogenetic stimulation of aIC terminals in the dorsolateral striatum induced larger excitatory post-synaptic currents and a smaller ratio of AMPA/NMDA currents recorded in medium spiny neurons compared to water-drinking mice (61). Interestingly, this synaptic plasticity was observed in male but not in female mice (61). Additionally, in mice, short-term ethanol exposure increases the excitability of pIC projection neurons targeting the bed nucleus of the stria terminalis neurons in females, but not males (62). Although the literature pinpoints the insular cortex as a key player in alcohol-related behaviors, sex differences in aIC and pIC function in binge and/or persistent alcohol intake despite quinine aversion remain to be uncovered.

Using the DID model, we repeatedly demonstrated a higher binge and aversion-resistant ethanol consumption in female compared to male mice. Combining the DID protocol with *in vivo* calcium fiber photometry recordings, we identified that aIC excitatory neurons are activated during the consumption of ethanol, ethanol adulterated with quinine, and water independently of sex. In the pIC, excitatory neurons are activated during binge ethanol drinking in both sexes, and during ethanol+quinine and water intake in females only. Importantly, sex differences during drinking behavior were identified only during ethanol+quinine consumption, with pIC neurons showing higher response in females than males. Then, using chemogenetics, we showed that inhibition of aIC glutamatergic neurons reduced bitter solution intake independently of solvent (e.g., ethanol or water) and sex. In contrast, inhibition of pIC glutamatergic neurons decreased the intake of quinine only in males and reduced persistent ethanol drinking exclusively in female mice.

Altogether, we identified a sex-dependent function of the pIC in persistent ethanol drinking, providing key elements to the characterization of insular cortex function in AUD in a sex-specific manner.

## METHODS AND MATERIALS

For additional details on the methods, see the Supplementary information.

### Stereotactic surgeries

#### Chemogenetic viral injection

The adeno-associated viruses encoding the inhibitory hM4Di receptor coupled to mCherry under the control of CaMKII promoter (AAV9-CaMKIIa-hM4D(Gi)-mCherry, titer >1.0e^+13^ vg/mL, Addgene, Watertown, Massachusetts, USA), to target glutamatergic neurons, or the control viral vector (AAV9/2-mCaMKII-mCherry-WPRE, titer 6.3e^+12^ vg/mL, ETH Zürich, Switzerland) were injected bilaterally (200-250 nL, 1 nL/sec) in the aIC (anteroposterior +1.7 mm; mediolateral ±3.1 mm; dorsoventral -3.5 mm from the bregma) or the pIC (anteroposterior -0.35 mm; mediolateral ±4.0 mm; dorsoventral -4.2 mm from the bregma).

#### Calcium fiber photometry viral injection and fiber implantation

A viral vector coding for the calcium sensor GCaMP6f (AAV9-CaMKII-GCaMP6f-WPRE-SV40, titer 2.5e^+12^ vg/mL, Addgene) was injected unilaterally (250 nL, 1 nL/sec) in the right aIC or pIC. An optic fiber (400 µm diameter, 0.48 NA, >90% efficiency) inserted in a ceramic ferrule was implanted 50 µm above the virus injection site.

### Drinking in the dark procedure (DID)

The DID protocol is based on the circadian rhythm where animals have access to different liquids (e.g., ethanol) during the nocturnal period (9,10). Two and a half hours after the onset of the dark phase, the water bottle in the home cage was replaced by different liquids for 2 hours per day for 4 consecutive days, followed by 3 days with water access only. This 7-day cycle is repeated for 4 weeks, starting with tap water (cycle 0), then ethanol (20% v/v in tap water, cycles 1 and 2) to study binge drinking behavior, and ethanol adulterated with quinine (500 µM of quinine, cycle 3) to study persistent drinking despite aversion. Finally, water or water adulterated with quinine (500 µM quinine) consumption was measured during a single 2-hour session and served as a control. Additionally, a subgroup of animals underwent three more days of water+quinine, followed by a last cycle of ethanol+quinine to control the resumption of persistent ethanol drinking (**Fig. 2A**). Daily liquid consumption was obtained by weighing the bottle before and after the 2-hour session, and mice were weighed on the first day of each cycle. Food was available *ad libitum* during drinking sessions performed in the home cages, and until the beginning of the sessions in multifunctional boxes for fiber photometry.

#### Blood ethanol concentration measurement

To measure blood ethanol concentrations (BEC), blood sampling was performed by incision of the lateral tail vein 30 minutes or 2 hours after the onset of the ethanol session. The samples were immediately centrifuged at 10,000 rpm for 10 minutes at 4°C. The BEC in mg/dL was obtained from the serum using the Analox GL5 analyzer. This experiment has been performed twice, and consistent results were observed between experiments. We excluded 3 males and 3 females from the BEC analysis after a 2-hour ethanol intake because the blood sample did not contain enough serum.

### Coding properties of pIC excitatory neurons during binge and persistent ethanol drinking

The same DID procedure as described before was used, except that no initial water cycle was performed. To study the coding properties of aIC and pIC neurons during binge and persistent ethanol drinking, the calcium signal and the drinking behavior (e.g., licking) have to be synchronized. Thus, after two ethanol binge cycles in their home cage, mice (n=30 males and n=31 females) underwent three cycles of ethanol drinking (cycles 3 to 5) in multifunctional boxes equipped with lickometers (**Fig. S2A**). Then, after one cycle of ethanol+quinine drinking in the home cages (cycle 6), mice underwent a cycle of ethanol+quinine drinking in the multifunctional boxes (cycle 7). Finally, a water and water+quinine photometry recording session was performed in these boxes as a control. The analysis of pIC neuronal activity during water and water+quinine consumption includes 3 males and 2 females, and 4 males and 2 females, respectively, who were ethanol-naive. The pIC recording experiment has been performed six times, aIC recording experiment has been repeated twice (**Table S2**). Replicated experiments showed consistent results.

#### Fiber photometry recordings

The recordings were performed with Neurophotometrics fiber photometry system (FP3002 V2, San Diego, California, USA), on the last day of the last ethanol and ethanol+quinine cycles, and during a single water and water+quinine session. The recordings were performed using a 4-branch patch cord (400 µm diameter, 0.48 NA, 1 branch per animal, Doric Lenses). Then, the drinking sprout was filled with the appropriate solution, and the recording lasted 1 hour as previously done (36). Mice were kept in the boxes for an additional hour for a total of a 2-hour session. The drinking sprout was automatically refilled to give *ad-libitum* access to the mice. The optimal wavelength for GCaMP6f excitation is 470 nm, while the isosbestic wavelength is 415 nm. To minimize the photobleaching effect of the recording, the light intensity at the tip of the patch cord was adjusted between 80 and 100 µW for both channels. The sampling rate was settled at 30 Hz for photometry recordings.

#### Calcium fiber photometry data analysis

Photometry recordings were analyzed using custom Python scripts https://github.com/praneethnamburi/Photometry_and_Licks_BeyelerLab. The timestamps for lick events were extracted and classified in bouts. To minimize potential confounds in the neuronal activity analyzed due to differences in the behavior across mice (e.g., number of licking bouts), the analysis of aIC and pIC neuronal activity was first performed on single licks and then on the combination of single licks and bouts (**Table S4, S5**), with a maximum of 5 licks/bouts per animal for both analyses. A bout is defined as a minimum of two licks with an inter-lick interval (ILI) less than or equal to 10 seconds for the peri-event analysis. The first 10 minutes of the test mice did not have access to liquid solutions, thus the signals were discarded from the analysis. Then, GCaMP and isosbestic signals were de-trended using the airPLS algorithm (63), and the detrended isosbestic signal was regressed from the detrended GCaMP signal. The resulting signal was bandpass filtered with cutoff frequencies of 0.2 and 6 Hz using a finite impulse response filter implemented in the scipy package. Finally, a time window of -10 to +10 s was defined for the peri-event analysis of the resulting signal aligned to each lick (time 0 s). We controlled that no event happened during the -10 to 0 s. The signal from each window was z-normalized by subtracting the mean of the baseline and dividing the result by the standard deviation of the baseline, where the baseline signal was defined as the signal from -10 s to -8 s before a lick onset. Z-normalized calcium signals were averaged for each male and female mouse during the reference (−8 to -5 s), the pre-lick (−3 to 0 s), the post-lick (0 to +3 s) and the post-ingestive (+6 to +9 s) windows to describe the signal changes.

#### Code availability

Photometry recordings were analyzed using custom Python scripts, which can be downloaded at: https://github.com/praneethnamburi/Photometry_and_Licks_BeyelerLab, no restrictions apply.

### Chemogenetic inhibition of aIC or pIC excitatory neurons during binge and persistent ethanol drinking

The same DID procedure as described above was used. The day before the chemogenetic manipulation an intraperitoneal (i.p.) injection of vehicle (NaCl, 0.1 mL/kg) was performed 30 minutes before the drinking session for habituation. The causal role of aIC or pIC glutamatergic neurons on drinking was tested after injection of CNO (i.p. 3 mg/kg in NaCl, Tocris, Bristol, United-Kingdom) in hM4Di mice 30 minutes before the last session of binge (day 18), persistent ethanol drinking (day 25) and a single session of water or water+quinine (day 29). To control for potential non-specific effects of CNO, mice expressing the control viral vector received either CNO or vehicle injections. As no behavioral differences were observed between the control-CNO and control-vehicle groups, these groups were pooled into a single control group. To note, some of the mice tested for chemogenetic inhibition of pIC on water+quinine drinking (day 29) were CNO naive. This experiment has been performed two times for the inhibition of pIC neurons and two times for the aIC inhibition (**Table S3**), with consistent results between experiment.

## RESULTS

### Behavioral characterization of binge ethanol consumption using the drinking in the dark protocol in male and female mice

After two 4-day cycles of ethanol consumption (**Fig. 1A**), the average 2-hour ethanol intake reported to the animal bodyweight, was higher in females than males (**Fig. 1B**, unpaired Student t-test, t=3.870, p=0.0007), whereas the blood ethanol concentration (BEC) was similar between sexes (**Fig. 1C**). After two hours of ethanol consumption, similar percentages of males (23%) and females (15%) reached the binge intoxication threshold of 80 mg/dL (Fischer’s exact test, p>0.9999). As previous studies suggest that mice drink most of the ethanol at the beginning of the DID sessions (11), we measured the proportion of ethanol consumed during the first 30 minutes of the 2-hour session. The proportion of ethanol consumed during the first 30 minutes was higher than the proportion drunk during the remaining 90 minutes for both males (67% vs 33%) and females (84% vs 16%), and was higher in females than males (84% vs 67%), supporting a pattern of binge consumption in both sexes, even stronger in females (**Fig. 1D**, 2-way ANOVA, sex factor F_1, 60_=0.000, p>0.999, ethanol proportion F_1, 60_=132.2, p<0.0001, with an interaction F_1, 60_=14.16, p=0.0004). Moreover, in an additional session, both ethanol intake and BEC were measured after 30 minutes of ethanol access, and were found to be higher in females compared to males (**Fig. 1E**, unpaired Student t-test, t=3.668, p=0.0009, and **Fig. 1F**, unpaired Mann-Whitney, U=71, p=0.0318 respectively). At this short time point, 75% of female and 37.5% of male mice reached the intoxication threshold of 80 mg/dL (Fisher’s exact test, p=0.0732).

**Fig. 1.**
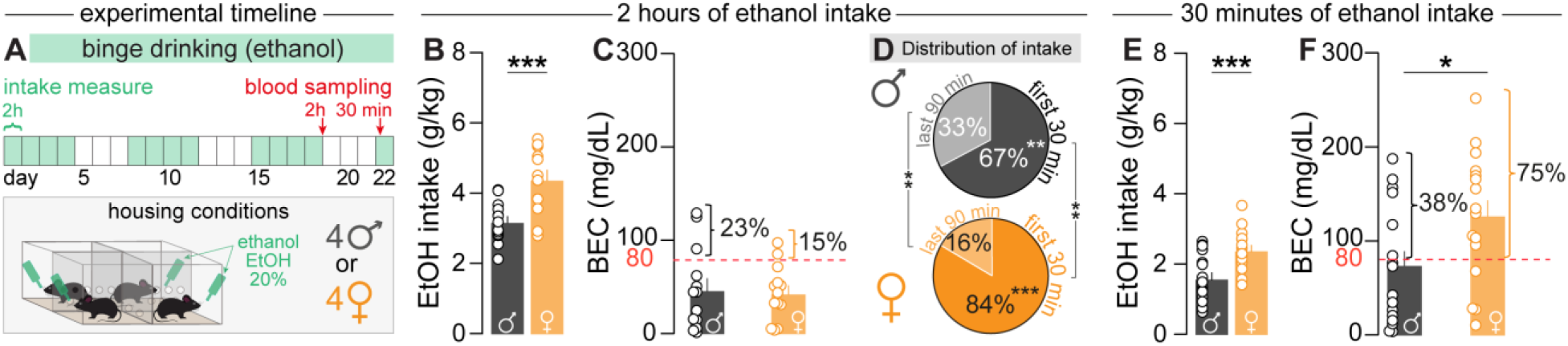
Validation of the drinking in the dark model. **A** Experimental timeline of binge ethanol exposure according to the drinking in the dark procedure. Blood was sampled after 2 hours of session on day 18 and after 30 minutes of session on day 22. **B, C** Average of ethanol intake (B) and blood ethanol concentration (C) after 2 hours of ethanol binge drinking (day 18) in male (n=13) and female (n=13) mice. The red dashed line represents the intoxication threshold of 80 mg/dL. **D** Proportion of ethanol intake during the first 30 minutes and last 90 minutes of a 2-hour session in male (n=16) and female (n=16) mice. **E, F** Average of ethanol intake (E) and blood ethanol concentration (F) after 30 minutes of session (day 22) in male (n=16) and female (n=16) mice. Data are shown as mean ± SEM. *p<0.05 **p<0.01 ***p<0.001 represent significant Student t-test or Mann-Whitney test.

### Sex differences in ethanol binge drinking during the drinking in the dark protocol

First, we analyzed drinking behavior across all sessions excluding the days of the chemogenetic test and independently of the viral vector injection site (e.g. aIC or pIC, **Fig. 2A**). Altogether, female mice had a higher level of water (**Fig. 2B**, 2-way RM-ANOVA, sex factor F_1, 109_=7.208, p=0.0084, session factor F_3, 327_=3.342, p=0.0195, without interaction F_3, 327_=0.1697, p=0.9168), ethanol (**Fig. 2D**, 2-way RM-ANOVA, sex factor F_1, 109_=116.7, p<0.0001, session factor F_5.331, 581.7_=0.0027, p=0.0027, without interaction F_6, 654_=0.9494, p=0.4589), and ethanol+quinine (**Fig. 2F**, 2-way RM-ANOVA, sex factor F_1, 108_=103.4, p<0.0001, session factor F_2, 216_=1.951, p=0.1446, without interaction F_2, 216_=1.951, p=0.1446) intake compared to males, across the sessions. This higher liquid intake in females was also observed on the average intake for the three solutions (**Fig. 2C**, unpaired Student t-test, t=2.685, p=0.0084, **Fig. 2E**, unpaired Student t-test, t=10.80, p<0.0001, **Fig. 2G**, unpaired Student t-test, t=10.17, p<0.0001). However, it is important to note that the difference between females and males is larger for ethanol as well as ethanol+quinine, compared to water (effect size Cohen’s d=0.04 for water 0.30 for ethanol and 0.25 for ethanol+quinine). Water, ethanol, and ethanol+quinine intake were higher in females than in males when expressed both in absolute volume (mL) and body weight-normalized volume (mL/kg, **Supplementary Fig. S1A-F**). To further analyze the link between binge ethanol drinking and persistent ethanol consumption, we correlated the average ethanol intake with the average ethanol+quinine intake. For male mice, we observed a positive correlation between binge drinking and persistent drinking (**Supplementary Fig. S1G**), and a strong tendency for a positive correlation in females (**Supplementary Fig. S1H**).

**Fig. 2.**
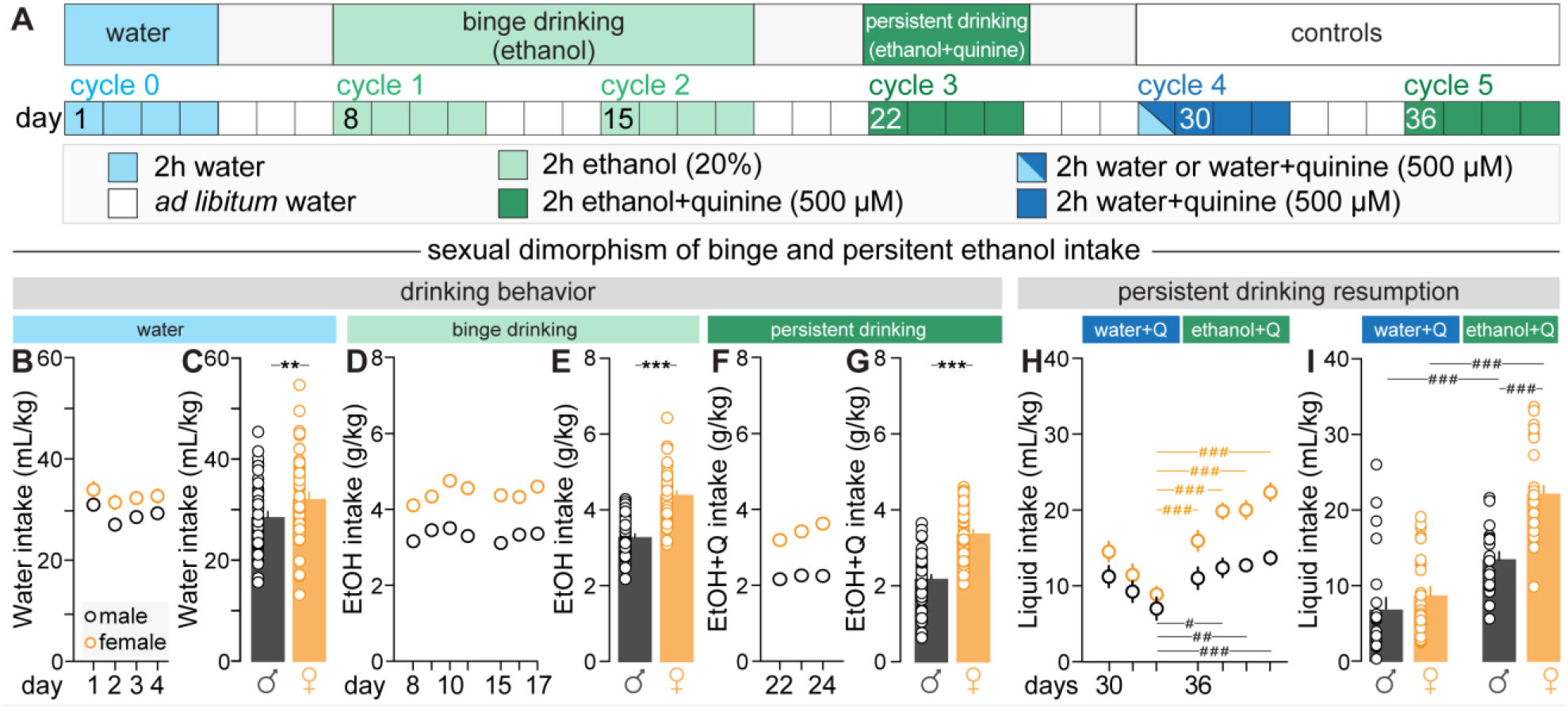
Sex differences in binge and persistent ethanol drinking. **A** Behavioral timeline during the drinking in the dark procedure. **B, C** Kinetic (B) and average (C) of water intake during cycle 0 in male (n=55) and female (n=56). **D, E** Kinetic (D) and average (E) of ethanol intake during cycles 1 and 2 in male (n=55) and female (n=56). **F, G** Kinetic (F) and average (G) of ethanol+quinine intake during cycle 3 in male (n=55) and female (n=55). **H** Kinetic of water+quinine (water+Q) and ethanol+quinine (ethanol+Q) intake in males (n=22) and females (n=26). **I** Comparison of water+quinine and ethanol+quinine intake during the last session of exposure in male (n=22) and female (n=26) mice. Data are shown as mean ± SEM. **p<0.01, ***p<0.001 represents a significant Student t-test. #p<0.05 ##p<0.01 ###p<0.001 represents a significant Bonferroni post-hoc test.

Finally, consumption of both ethanol and ethanol adulterated with quinine was independent of the hormonal cycle in female mice (**Supplementary Fig. S1I-K**). To test the strength of persistent ethanol drinking despite quinine aversion, a subgroup of mice that had undergone binge and persistent drinking had access to water+quinine only for 3 sessions, followed by 4 sessions of ethanol+quinine access. Interestingly, both males and females consume more ethanol+quinine than water+quinine (**Fig. 2H**, 2-way RM-ANOVA, sex factor F_1, 46_=20.64, p<0.0001, session factor F_6, 276_=23.41, p<0.0001, with an interaction F_6, 276_=3.450, p=0.0027). This observation was validated on average, as for both sexes, consumption of ethanol+quinine was larger than water+quinine (**Fig. 2I**, 2-way RM-ANOVA, sex factor F_1, 46_=17.85, p=0.0001, liquid factor F_1, 46_=85.86, p<0.0001, with an interaction F_1, 46_=9.787, p=0.0030). These results validate the appetitive property of ethanol, as its addition to quinine is sufficient to increase quinine drinking in both sexes. Therefore, we named the consumption of ethanol+quinine persistent drinking despite aversion.

### Neural activity of right aIC and pIC glutamatergic neurons during binge and persistent ethanol drinking

We used fiber photometry with the genetically encoded calcium sensor GCaMP6f (**Fig. 3A, S2A**), to record aIC (**Fig. 3B, S2B**) and pIC (**Fig.3C, S2K**) glutamatergic neuron activity during single lick of different solutions over 1-hour sessions (**Fig. 3D**). Behaviorally, the average ethanol and ethanol+quinine intake across all drinking sessions, excluding the recording session, was higher in females than males (**Fig. 3E**, unpaired Mann-Whitney test, U=145, p=0.0143, **Fig. 3F**, unpaired Student t-test, t=2.357, p=0.0233). For animals with neuronal recordings in the right aIC, a similar average number of single licks or events (i.e., combination of single licks and licking bouts) were analyzed across ethanol, ethanol+quinine, water, and water+quinine conditions (**Supplementary Fig. S2C-J**). Comparable results were obtained in animals recorded in the right pIC (**Supplementary Fig. S2L-O, R, S**), except for the water condition, in which males exhibited a higher number of single licks and events compared to females (**Supplementary Fig. S2P**, unpaired Student’s t-test, t=2.382, p=0.0309; **Supplementary Fig. S2Q**, unpaired Mann-Whitney test, U=119, p=0.03).

**Fig. 3.**
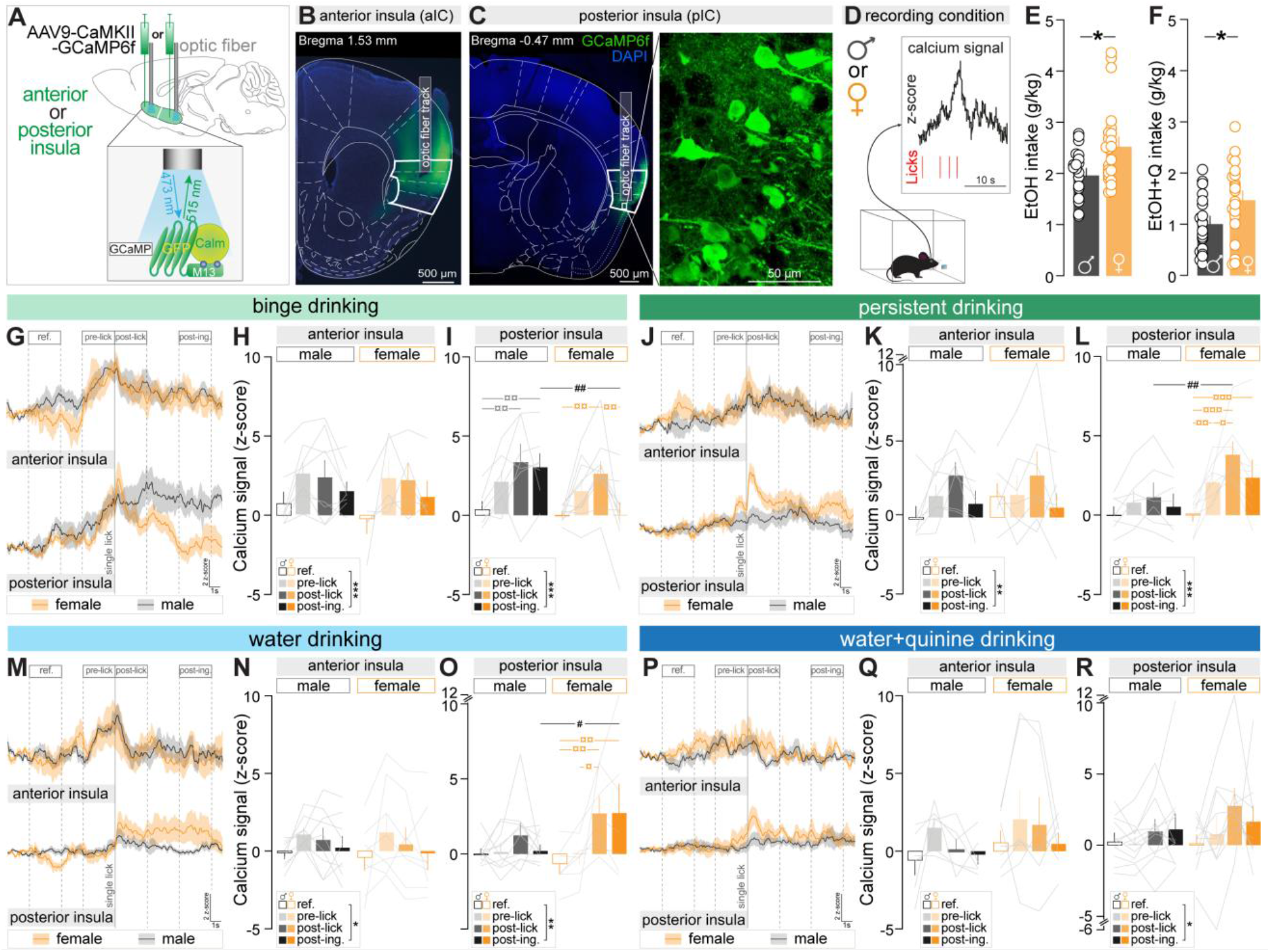
Coding properties of anterior (aIC) and posterior (pIC) insular cortex excitatory neurons during single licks of binge and persistent ethanol consumption. **A** Viral strategy to record calcium changes in aIC or pIC excitatory neurons. A viral vector carrying the gene coding for the calcium sensor GCaMP6f was injected unilaterally in the aIC or the pIC. **B, C** Representative image of GCaMP6f expression in aIC (B) or pIC (C) excitatory neurons. The mouse brain atlas delineation has been overlaid to the image, with the borders of the aIC or pIC in bold. **D** Conditions during photometry recordings to synchronize calcium signal and licking behavior. **E** Average ethanol intake during all binge-drinking sessions except the recording day in male (n=21) and female (n=24) mice. **F** Average ethanol+quinine intake during all persistent drinking sessions except the recording day in male (n=21) and female (n=22) mice. **G** Peri-ethanol licks analysis of the calcium signal in the aIC (up) and pIC (bottom) during single ethanol licks in male and female mice. **H, I** Average of calcium signal in the aIC (H) or pIC (I) during ethanol licking for the reference, pre-lick, post-lick, and post-ingestive periods in male (n=8 for aIC and n=7 for pIC) and female (n=5 for aIC and n=10 for pIC) mice. **J** Peri-ethanol+quinine licks analysis of the calcium signal in the aIC (up) and pIC (bottom) single ethanol+quinine licks in male and female mice. **K, L** Average of calcium signal in the aIC (K) or pIC (L) during ethanol+quinine licking for the reference, pre-lick, post-lick, and post-ingestive periods in male (n=7 for aIC and n=7 for pIC) and female (n=7 for aIC and n=8 for pIC) mice. **M** Peri-water licks analysis of the calcium signal in the aIC (up) and pIC (bottom) during single water licks in male and female mice. **N, O** Average of calcium signal in the aIC (N) or pIC (O) during water licking for the reference, pre-lick, post-lick, and post-ingestive periods in male (n=9 for aIC and n=10 for pIC) and female (n=8 for aIC and n=7 for pIC) mice. **P** Peri-water+quinine licks analysis of the calcium signal in the aIC (up) and pIC (bottom) during single water+quinine licks in male and female mice. **Q, R** Average of calcium signal in the aIC (Q) or pIC (R) during water+quinine licking for the reference, pre-lick, post-lick, and post-ingestive periods in male (n=6 for aIC and n=12 for pIC) and female (n=7 for aIC and n=10 for pIC) mice. Data are shown as mean ± SEM. *p<0.05 **p<0.01 ***p<0.001 represent significant Student t-test, or Mann-Whitney test, or a main effect of the 2-way RM ANOVA. #p<0.05, ##p<0.01, ¤p<0.05, ¤¤p<0.01, ¤¤¤p<0.01 represent significant Bonferroni post-hoc tests.

At the neural level, we segmented the signal recorded in the insular cortex of the right hemisphere in four 3-seconds time windows centered around licking onset: reference (−8 to -5 s), pre-lick (−3 to 0 s), post-lick (0 to +3 s), and post-ingestive (+6 to +9 s, **Fig. 3**). When centered around ethanol consumption, the activity of aIC neurons was different across the four windows, independently of sex (**Fig. 3G, H**, 2-way RM-ANOVA, sex factor F_1, 11_=0.2616, p=0.6191, period factor F_3, 33_=7.374, p=0.0007, without interaction F_3, 33_=0.3421, p=0.7950). In contrast, the activity of pIC neurons was increased during ethanol post-lick compared to the reference window, in both males and females, and was higher in males during the post-ingestive window, compare to the male reference and to the female post-ingestive windows (**Fig. 3I**, 2-way RM-ANOVA, sex factor F_1, 15_=2.916, p=0.1083, period factor F_3, 45_=9.974, p<0.0001, with an interaction F_3, 45_=2.875, p=0.0465).

Regarding ethanol+quinine drinking, aIC neurons activity was different across periods, independently of sex (**Fig. 3J, K**, 2-way RM-ANOVA, sex factor F_1, 12_=0.1030, p=0.7538, period factor F_3, 36_=4.757, p=0.0068, without interaction F_3, 36_=0.7980, p=0.5031). Interestingly, pIC neuronal activity around ethanol+quinine licking varied only in female mice. First, we observed higher pIC neuron activation during the pre-lick period compared to the reference, indicating an anticipatory activation of these neurons. Neuronal activity was further increased during the post-lick period compared to pre-lick, indicating that the peak activity response is occurring triggered by ethanol+quinine licking. Notably, pIC neural activity was higher in females than in males, specifically during this ethanol+quinine post-lick period, when licking occurred (**Fig. 3L**, 2-way RM-ANOVA, sex factor F_1, 13_=3.447, p=0.0862, period factor F_3, 39_=11.75, p<0.0001, with an interaction F_3, 39_=3.717, p=0.0192).

Additionally, we performed fiber photometry recordings during water and water+quinine licking. The activity of aIC neurons was different across periods, independently of sex (**Fig. 3M, N**, 2-way RM-ANOVA, sex factor F_1, 15_=0.09366, p=0.7638, period factor F_3, 45_=3.956, p=0.0138, without interaction F_3, 45_=0.1703, p=0.9159), whereas pIC neuron activity was increased in females during post-lick and post-ingestive periods relative to the reference, and was higher in females than males in the post-ingestive period (**Fig. 3O**, 2-way RM-ANOVA, sex factor F_1, 15_=1.101, p=0.3107, period factor F_3, 45_=6.288, p=0.0012, with an interaction F_3, 45_=2.814, p=0.0498). For water+quinine licking, the activity of aIC neuron was similar across periods, while it was different for pIC neurons, independently of sex (**Fig. 3P-R**, 2-way RM-ANOVA for pIC, sex factor F_1, 20_=0.9419, p=0.3434, period factor F_3, 60_=2.962, p=0.0392, without interaction F_3, 60_=0.8110, p=0.4928). Similar patterns of neuronal activity were observed in the aIC when analyzing the first 5 events (i.e., combination of single licks and licking bouts, **Supplementary Fig. S3A-B, S3D-E, S3G-H, S3J-K**). In the pIC, comparable neuronal activity was observed when analyzing the first 5 events for all liquids (**Supplementary Fig. S3F, S3I, S3L**), except for ethanol where pIC activity was different across periods, without specific effects (**Supplementary Fig. S3C**, 2-way RM-ANOVA, sex factor F_1, 16_=0.6862, p=0.4196, period factor F_3, 48_=13.06, p<0.0001, without interaction F_3, 48_=0.5825, p=0.6294).

Furthermore, we assessed whether consumption history calculated as the average ethanol or ethanol+quinine intake across all sessions except the recording session, correlated with aIC and pIC neuronal calcium signals during consumption. Activity of aIC neurons did not correlate with intake history in either sex (**Supplementary Fig. S4A-D**). Similarly, pIC activity showed no correlation with ethanol intake history in males or females (**Supplementary Fig. S4E, F**), nor with ethanol+quinine intake in males (**Supplementary Fig. S4G**). However, females exhibited a trend toward a positive correlation between ethanol+quinine intake history and pIC activity (**Supplementary Fig. S4H**, Pearson’s R^2^=0.46, p=0.07).

### Chemogenetic inhibition of aIC and pIC glutamatergic neurons during binge and persistent ethanol drinking

To study the causal role of the aIC and pIC excitatory neurons in binge ethanol drinking and persistent ethanol drinking (ethanol adulterated with quinine), we used a chemogenetic approach (**Fig. 4A-C**) to inhibit aIC or pIC glutamatergic neurons by expressing hM4Di, under the control of the CaMKIIɑ promoter. Beforehand, we tested the viral approach using *in-vivo* electrophysiology and confirmed a decrease in the average firing rate from 30 to 55 minutes after CNO injection compared to baseline (**Supplementary Fig. S4A, B**, paired Wilcoxon test, p=0.0232). Chemogenetic inhibition of aIC glutamatergic neurons (**Supplementary Fig. S5C**) did not influence ethanol drinking in both sexes (**Fig. 4D**). The same neuronal manipulation reduced ethanol+quinine drinking independently of sex (**Fig. 4E**, 2-way ANOVA, sex factor F_1, 48_=15.78, p=0.0002, virus factor F_1, 48_=6.180, p=0.0165, interaction F_1, 48_=3.151, p=0.0822). Moreover, inhibition of aIC neurons did not change water consumption in males or females (**Fig. 4F**), whereas it decreased water+quinine drinking independently of the sex (**Fig. 4G**, 2-way ANOVA, sex factor F_1, 25_=1.541, p=0.2259, virus factor F_1, 25_=14.72, p=0.0008, interaction F_1, 25_=3.839, p=0.0613).

**Fig. 4.**
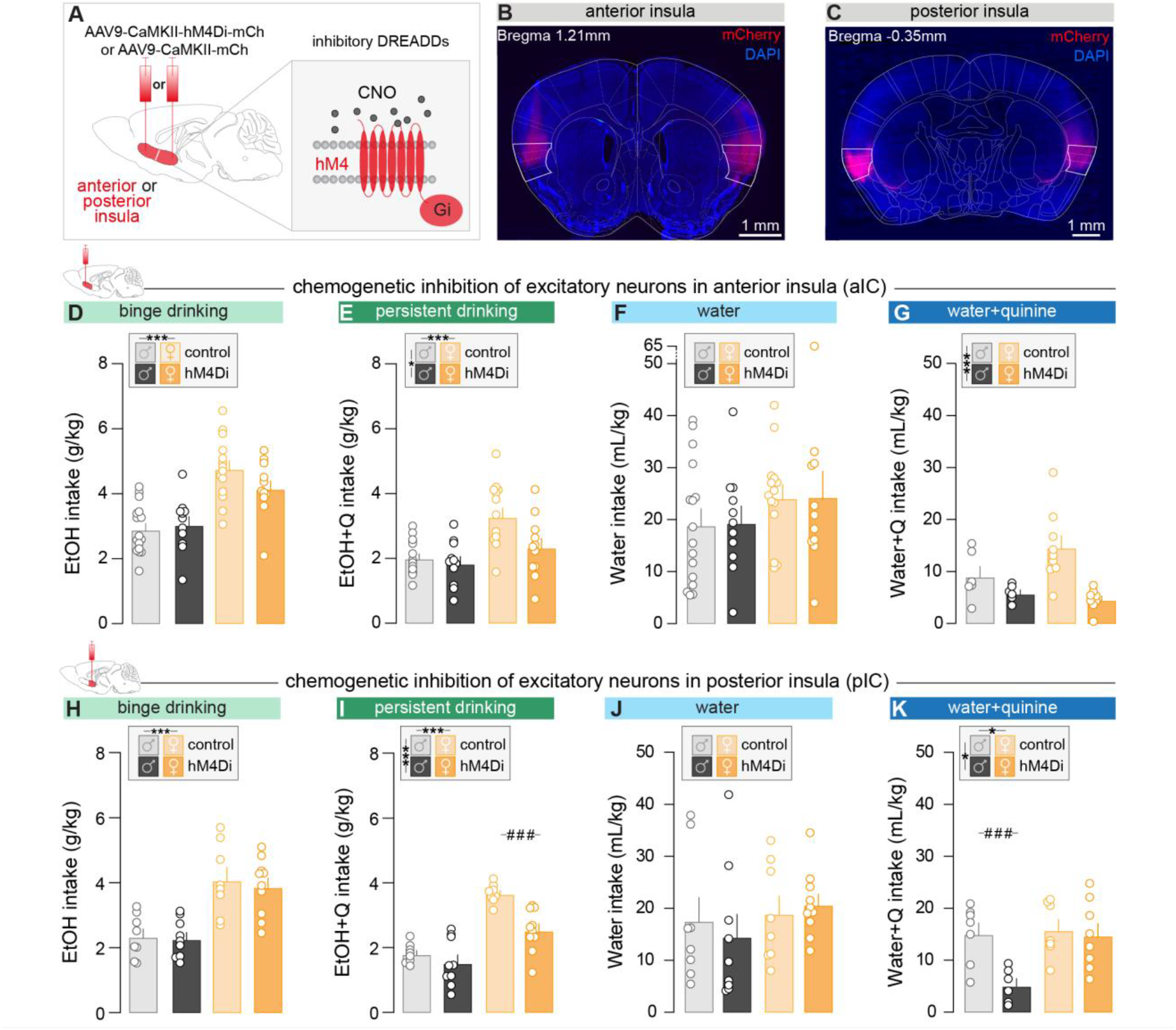
Chemogenetic inhibition of anterior (aIC) or posterior (pIC) insular cortex excitatory neurons during binge and persistent ethanol intake. **A** Experimental design to inhibit aIC and pIC excitatory neurons. A viral vector carrying the gene coding for the inhibitory receptor hM4Di fused to the fluorescent reporter mCherry, or the control virus, containing only the gene for mCherry, was bilaterally injected in the aIC or pIC. **B, C** Representative images of the chemogenetic viral vector expression in aIC (B) and pIC (C) neurons. The mouse brain atlas delineation has been overlaid on the image, with the borders of the aIC and pIC in bold. **D** Average ethanol intake during chemogenetic manipulation of aIC glutamatergic neurons (day 18) in male (control n=16, hM4Di n=11) and female (control n=14, hM4Di n=12) mice. **E** Average of ethanol+quinine intake during chemogenetic manipulation of aIC neurons (day 25), in male (control n=16, hM4Di n=11) and female (control n=14, hM4Di n=11) mice. **F** Average water intake during chemogenetic manipulation of aIC neurons (day 29), in male (control n=16, hM4Di n=11) and female (control n=14, hM4Di n=11) mice. **G** Average of water+quinine intake during chemogenetic manipulation of aIC neurons (day 29), in male (n=6 for both groups) and female (control n=9, hM4Di n=8) mice. **H** Average ethanol intake during chemogenetic manipulation of pIC neurons (day 18), in male (control n=8, hM4Di n=9) and female (control n=8, hM4Di n=10) mice. **I** Average of ethanol+quinine intake during chemogenetic manipulation of pIC neurons (day 25), in male (control n=8, hM4Di n=9) and female (control n=8, hM4Di n=10) mice. **J** Average of water intake during chemogenetic manipulation of pIC neurons (day 29), in male (control n=8, hM4Di n=9) and female (control n=8, hM4Di n=10) mice. **K** Average of water+quinine intake during chemogenetic manipulation of pIC neurons (day 29), in an independent group of males (control n=7, hM4Di n=6) and female (control n=7, hM4Di n=8) mice. Data are shown as mean ± SEM. *p<0.05 ***p<0.001 represents the significant main effect of the 2-way ANOVA. ###p<0.001 represents a significant Bonferroni post-hoc test.

Similarly, to the aIC, inhibition of pIC excitatory neurons (**Supplementary Fig. S5D**) did not influence ethanol drinking in both sexes (**Fig. 4H**). However, pIC inhibition reduced ethanol+quinine intake exclusively in female mice (**Fig. 4I**, 2-way ANOVA, sex factor F_1, 31_=58.57, p<0.0001, virus factor F_1, 31_=14.24, p=0.0007, with an interaction F_1, 31_=5.263, p=0.0287). As for aIC inhibition, pIC silencing also did not alter water consumption in males nor females (**Fig. 4J**). However, in contrary to aIC, pIC silencing altered water+quinine intake in male mice (**Fig. 4K**, 2-way ANOVA, sex factor F_1, 24_=6.284, p=0.0194, virus factor F_1, 24_=7.063, p=0.0138, with an interaction F_1, 24_=4.582, p=0.0427).

To rule out any behavioral confound due to the treatment, CNO or vehicle injection was performed in control mice. CNO injection in control animals did not alter ethanol, ethanol+quinine or water intake compared to control animals injected with vehicle (**Supplementary Fig. S5E-G, I-K**). Finally, we found that locomotion was similar between control mice injected with vehicle or CNO, and mice expressing the inhibitory DREADDs injected with CNO (**Supplementary Fig. S5H, L**). Together, our results demonstrate a sex-dependent function of the antero-posterior insula axis on drinking behaviors. The aIC neurons are recruited for bitter solution drinking independently of sex, whereas the pIC sustains persistent ethanol drinking in females, and water+quinine drinking in males.

## DISCUSSION

The goal of our study was to elucidate whether the aIC and pIC exhibit sex-dependent functions in ethanol binge drinking and ethanol persistent drinking despite bitter aversion in mice. First, we observed a higher binge and persistent ethanol intake in females compared to males (**Fig. 2E, G)**. Second, at the neuronal activity level, we showed that aIC neuronal activity was increased around ethanol, ethanol+quinine, and water across all periods, independently of sex (**Fig. 3H, K, N**), but not around water+quinine consumption (**Fig. 3Q**). In addition, right pIC neuronal activity increased in both sexes during ethanol drinking (**Fig. 3I**), and only in females during ethanol+quinine and water consumption (**Fig. 3L, O**). However, sex differences during licking behavior were observed only for ethanol+quinine drinking, where pIC neural activity was higher in females than in males. For water+quinine licking, we observed increased activity of pIC neurons during water+quinine licking across periods, independently of sex (**Fig. 3R**). Additionally, we demonstrated that chemogenetic inhibition of aIC glutamatergic neurons reduced the intake of quinine-adulterated solutions (e.g., ethanol and water, **Fig. 4E, G**) regardless of sex, whereas chemogenetic inhibition of pIC glutamatergic neurons reduced persistent ethanol drinking exclusively in females and quinine drinking in males (**Fig. 4I, K**). Altogether, these findings suggest a topographical and sex-dependent divergent function of the insular cortex in bitter liquid drinking, highlighting the pIC glutamatergic neurons as a substrate of persistent ethanol drinking in female mice.

### Sex-dependent behavior on binge and persistent ethanol drinking

Our results repeatedly showed a higher binge and persistent ethanol intake in females compared to males, independently of the estrous cycle (**Supplementary Fig. S1J, K**), which is consistent with and extends previous literature highlighting sex differences in drug addiction (64,65). Indeed, multiple preclinical studies also reported a higher binge (13,28,62) and persistent (13,29,66) ethanol intake per body mass in females compared to males. Importantly, to rule out any confound on sex-dependent aversion resistance for quinine, we adulterated the ethanol solution with 500 µM quinine, a concentration for which both male and female mice have similar aversive thresholds (67). Therefore, the sex difference observed in the consumption of ethanol adulterated with quinine is independent of the aversion sensitivity but driven by the motivation to consume ethanol. We further demonstrated that females exhibited higher (BEC) than males after 30 minutes of alcohol consumption (**Fig. 1F**). This difference may reflect sex-dependent variations in ethanol metabolism and absorption. Indeed, Robinson et al. (68) reported that, following intragastric administration of ethanol (1 g/kg), BEC peaked more rapidly in female than male rats, indicating faster gastric absorption in females. In addition, several studies have shown that females metabolize ethanol more rapidly than males (69,70), which supports our results showing that, after 2 hours of ethanol intake, a lower percentage of females reaches the BEC intoxication threshold of 80 mg/dL than males (**Fig. 1C**).

A key advantage of the DID model is that it rapidly induces high levels of ethanol drinking and intoxication in mice without food or water deprivation, similar to what is observed during human binge alcohol drinking. The protocol can also be adapted to model persistent drinking despite aversion by adding quinine to ethanol. This within-animal approach allows investigation of the transition from episodic binge drinking to persistent alcohol consumption. While quinine adulteration is widely used to mimic persistent ethanol drinking despite aversive consequences, it does not capture the broader social, legal, and health negative consequences associated with AUD in humans, nor its slow, multifactorial development. We acknowledge this limit in our study, and for this reason, we chose to refer to “persistent” rather than “compulsive” ethanol intake.

### Coding properties of right aIC glutamatergic neurons during binge and persistent ethanol intake

Our results revealed that right aIC excitatory neurons are activated during ethanol and ethanol+quinine consumption across periods independently of sex (**Fig. 3H, K**). A recent study demonstrated that the projection from the aIC to the dorsolateral striatum encodes binge ethanol consumption in males but not females (61). In contrast, our recordings measured the global glutamatergic neuronal population in the aIC, which may account for the differing results. These findings suggest that sex-dependent functions of the aIC during binge drinking may be mediated by distinct neural projections. For persistent ethanol consumption, few studies have investigated the aIC neuronal activity after ethanol+quinine drinking. Chen and Lasek reported an increased expression of the neuronal activity marker *cFos* in the aIC after ethanol+quinine intake compared to water, ethanol and quinine drinking in male mice (71). Additionally, a recent *in vivo* electrophysiological study in male rats demonstrated that aIC neurons with increased firing at ethanol drinking onset also enhanced their activity during ethanol+quinine consumption, demonstrating that aIC neurons in males code both binge and persistent ethanol drinking (72). However, to our knowledge, no study has examined the dynamics of aIC activation in females during persistent alcohol consumption. Thus, our results corroborate previous findings and confirm the role of the right aIC in persistent consumption in males, and extend this to females.

Finally, our results showed increased response of right aIC glutamatergic neurons around water consumption (**Fig. 3N**), but not water+quinine intake (**Fig. 3Q**). For water intake, this is in line with and extends the findings of a recent study highlighting that aIC neurons were activated after water licking in male mice (73). For water+quinine consumption, Chen and colleagues revealed aIC activation after quinine tasting (71). However, these findings were obtained using a two-bottle choice paradigm, in which quinine was presented alongside with water, allowing animals to choose between the two solutions. This procedural difference likely accounts for the discrepancy with our results, as the availability of a competing non-aversive solution engages specific choice and value-comparison circuits, which include the aIC, that is not recruited in our single-solution DID paradigm.

### Function of aIC excitatory neurons in bitter taste processing

Our results revealed that aIC neuron inhibition does not alter binge ethanol intake in both sexes. These results support and extend Haaranen et al. study, which showed that chemogenetic inhibition of aIC neurons did not affect alcohol intake in alcohol-preferring male rats in a two-bottle choice model (58). However, it contrasts with previous literature demonstrating that chemogenetic manipulation of aIC neurons modulates alcohol binge drinking, although the results are divergent. Indeed, while silencing aIC neurons increased operant responses for alcohol (41), it decreased alcohol intake in a two-bottle choice paradigm (74). Altogether, these results suggest that aIC function in ethanol binge drinking might be dependent on the animal model used. Moreover, while we specifically manipulated aIC glutamatergic neurons, the studies cited previously manipulated all aIC mature neurons independently of their phenotype, which could explain some discrepancy in the results observed.

We also demonstrated that inhibition of aIC glutamatergic neurons reduced the intake of solutions adulterated with quinine (e.g., ethanol or water) independently of sex. These results confirm and extend previous experiments performed in male rats and showing a reduction of ethanol+quinine intake after inhibition of aIC neurons, as well as inhibition of aIC projections to locus coeruleus or nucleus accumbens core (74,75). Thus, our results bring additional and complementary information to these studies.

Regarding the role of aIC in water+quinine intake, the literature is divergent and, to our knowledge, only addresses this question in males so far. On one hand, optogenetic inhibition or lesion of the aIC respectively had no effect (76,77) or decreased (78) sensitivity to quinine, leading to an increase in intake. On the other hand, activation of aIC neurons and their projection to the basolateral amygdala also decreased quinine sensitivity (47,79). As in our study aIC excitatory neuron inhibition had no effect on water+quinine intake, our findings are consistent with two studies (76,77) and contrast with three others (42, 28, 69). Moreover, we chose to not water-deprive our animals to observe more ethological behavioral responses. In contrast, in the other studies, animals were water-deprived before quinine exposure, which could likely have induced a physiological stress altering aIC response to quinine. In addition, Staszko et al., have shown that previous taste experience with water adulterated with quinine decreases the number of active cells and neuronal response in the aIC (80). Thus, in our study, the absence of modulation of water+quinine intake by aIC neuron inhibition could be explained by pre-exposure to quinine before chemogenetic manipulation. Finally, *cFos+* neuron distribution in aIC after quinine exposition significantly varies across aIC subareas and the median-lateral axis (81), whereas in our study we targeted the entire aIC. Thus, further investigations are necessary to sharpen our understanding of aIC neurons in aversive solution consumption in both sexes.

### Function of the insular cortex in processing information from the peripheral nervous system

The insular cortex is a key brain region integrating taste perception and post-ingestive effects of drug intake. For example, Naqvi and Bechara (82), proposed that the insular cortex integrates the conscious pleasure generated by the bodily effect of drug intake and the pain following withdrawal. In the context of AUD, alcohol taste activates gustatory pathways, including the insula (83), and a recent study demonstrated a positive correlation between aversive taste reactivity to orally delivered 20% EtOH and pIC cFos+ cells in rats (84). In addition, fMRI studies revealed that the insular cortex is activated after gastric distention in rodents (85,86), and that the pIC in particular receives interoceptive visceral information (87). However, to our knowledge, no studies have investigated how peripheral information after alcohol ingestion is integrated by the insula to regulate subsequent alcohol consumption in rodents. Additional studies are necessary to understand how the integration of gustatory and post-ingestive signals by the insula shapes alcohol consumption in both sexes.

### Functional sex differences in right pIC coding properties of persistent ethanol drinking

To decipher how pIC excitatory neurons encode binge drinking and persistent ethanol drinking, we performed *in vivo* fiber photometry recordings of the right pIC. We observed increased activity of right pIC glutamatergic neurons in response to ethanol in both sexes (**Fig. 3I**). In contrast, activity increased only in females for ethanol+quinine and water (**Fig. 3L, O**). Importantly, sex differences during the licking period itself are detected exclusively during ethanol+quinine drinking with females showing higher right pIC neuron activity than males (**Fig. 3L**). In contrast, during ethanol and water consumption, sex differences emerged after the licking behavior, within the post-ingestive period (+6 to +9 s). Finally, during water+quinine drinking, pIC neuronal activity was different across periods, independently of sex (**Fig. 3R**).

Our results showed a sex-dependent functional dynamic of right pIC neuronal activity in persistent ethanol drinking (**Fig. 3L**). In physiological conditions, Lezzi et al. showed that pIC neurons in females are less excitable than in males (89). In contrast, a recent study demonstrated that short ethanol exposure increases the intrinsic excitability of pIC neurons projecting to the bed nucleus of the stria terminalis, in female mice only (62). This study, strengthened by our present results, suggests that ethanol consumption modulates pIC neuronal excitability. Altogether, it highlights pIC glutamatergic neurons and their projections as a key neural population supporting sex-dependent phenotypes in alcohol-related behaviors. Moreover, we revealed sex differences in pIC neuronal activity in the post-ingestive period after ethanol and water consumption, which was higher in males for ethanol, and higher in female for water (**Fig. 3I, O**). These results may reflect a sex-specific processing of post-ingestive interoceptive signals by the pIC, rather than differences strictly related to the consummatory behavior. Finally, right pIC neuronal activity increased during water+quinine drinking, which is consistent with previous studies reporting that pIC neurons encode bitter taste, independently of the sex (49,88).

Importantly, some literature suggests a lateralization of insular cortex functions, including water and alcohol drinking. Indeed, several neuroimaging studies report lateralized right-sided posterior insula atrophy in AUD patients (56,90,91). For this reason, we focused fiber photometry recordings on the right insula. However, unilateral recordings remain one of the limitations of our study. Understanding the lateralization of insular cortex function in alcohol consumption is an important question that will require further investigation.

### Uncovering sex-dependent pIC function in persistent ethanol intake

Fiber photometry recordings reflect the functional dynamics of pIC glutamatergic neurons in response to ethanol intake, however, it does not provide information on its causal role on persistent ethanol intake. To address this question, we used chemogenetic manipulations to demonstrate the necessity of pIC excitatory neurons to change persistent ethanol drinking in females. We demonstrated sex differences in the function of pIC glutamatergic neurons in drinking solutions adulterated with bitter taste. To our knowledge, these results are the first demonstration of a sex-dependent function of pIC neurons on persistent ethanol drinking. Indeed, in males, inhibition of pIC glutamatergic neurons reduced only water adulterated with quinine intake (**Fig. 4K**), while in females, the same neuronal manipulation decreased exclusively persistent ethanol drinking (**Fig. 4I**). Previous literature highlighted the role of the insular cortex in aversive taste processing (88,92). While the function in bitter tastant has been first dedicated to the pIC (47), multiple studies revealed that the distribution of neurons activated by quinine is not strictly specific to the pIC and can be found in the entire insular cortex (73–76). However, so far, no study has investigated the causal role of the insular cortex in the anteroposterior axis for bitter-tastant solution drinking. Therefore, our results provide new evidence on how the insular cortex influences bitter-solutions drinking in a sex-dependent manner. Using the two-bottle choice paradigm, a recent study showed a higher expression of the neuronal activity marker *cFos* in the pIC of males compared to females after ethanol+quinine intake (93). Importantly, in this study ethanol was adulterated with 100 µM of quinine. A previous report suggests that females are not sensitive to ethanol adulterated with quinine at this concentration whereas males are (29) which could explain the discrepancy with our results where we used a concentration of quinine at 500 µM. Furthermore, using an operant alcohol drinking model in male rats, pharmacological inactivation of pIC neurons reduces alcohol intake (59) while in our study, chemogenetic inhibition of pIC excitatory neurons did not change ethanol drinking nor in males neither in females. Several methodological aspects can, at least in part, support this discrepancy. First, operant and voluntary alcohol drinking could recruit different neuronal circuits, second, the methods used for the neuronal inhibition (e.g. pharmacology vs chemogenetics) and the animal model used (e.g. rat vs mouse) were different.

To rule out the possibility that pIC glutamatergic inhibition was due to the non-selective effects of CNO, we performed multiple control experiments. Indeed, it has been shown that CNO can bind to non-DREADD targets, and does not cross the blood-brain barrier. However, clozapine, a CNO metabolite, is reaching the brain and can also have multiple pharmacological targets that can induce several physiological and behavioral effects (94,95). In our control experiments, we found that CNO injection in mice expressing the control viral vector in pIC had no effects on either liquid consumption (e.g., ethanol, ethanol+quinine or water) or locomotion. Furthermore, CNO injection in mice injected with the viral vector carrying the gene coding for hM4Di also did not affect locomotion. These results indicate that CNO does not have non-specific effects either on drinking or locomotion and secondly, that the inhibition of pIC excitatory neurons did not affect locomotion. Overall, these results confirm that the behavioral effect of CNO on ethanol+quinine drinking in females and water+quinine intake in males is due to selective inhibition of pIC excitatory neurons. Finally, in our study the manipulation and the recording of pIC excitatory neurons was performed regardless of their outputs. However, the pIC is a highly wired region sending dense projections to several regions (49,96), including the central amygdala, a brain region highly involved in alcohol-related behaviors and supporting sex differences in alcohol intake (97–99). Therefore, future investigations will be crucial to understand the sex-dependent function of pIC neurons in alcohol-related behaviors at the circuit level.

### Theoretical circuit mechanism within the pIC underlying sex differences in persistent ethanol drinking in mice

Persistent ethanol drinking typically reflects a behavior in which reward and aversion counterbalance each other, driving the pursuit of the reward (e.g., ethanol) despite the associated aversion (e.g., quinine). In our study, female mice drink more ethanol+quinine than males (**Fig. 2G**), indicating that the balance between both modalities is shifted toward the reward. At the neuronal level, we demonstrated that this sex difference engages pIC glutamatergic neurons (**Fig. 3L, 4I**). Interestingly, the literature reports that pIC neurons can mediate both reward and aversive behavior in a projection-specific manner. Indeed, optogenetic activation of pIC to the central amygdala (CeA) induces avoidance behavior and learning in male and female mice (100), showing that this pathway drives aversive-like behaviors in both sexes. In contrast, using a nosepoke-triggered photostimulation paradigm or a real-time place preference task, Girven and colleagues showed that photostimulation of pIC to the bed nucleus of the stria termianlis (BNST) terminals is rewarding in both male and female mice (101), demonstrating that this pIC output mediates reward-related behaviors. Thus, in our experimental context, both pIC-CeA and pIC-BNST projections may be, respectively, engaged by the aversive and rewarding components of ethanol+quinine. Moreover, ethanol drinking selectively enhances the excitatory properties of pIC to BNST neurons in females but not males (62). Consequently, ethanol drinking may potentiate the reward-associated pIC-BNST pathway in females, shifting circuit engagement toward reward over the pIC-CeA aversion circuit, thus promoting higher persistent ethanol drinking in females than males. Finally, this mechanism may additionally reflect modulation through local GABAergic microcircuitry in the pIC (**Supplementary Fig. S6**).

### Translation relevance

Human studies have primarily examined neurobiological alterations of the aIC in AUD men and women, reporting reduced gray matter density compared to healthy controls (50–53). In addition, AUD men and women had reduced aIC volume compared to healthy controls (50), which is associated with higher compulsivity and impulsivity. In our study, we showed that aIC encodes and is required for persistent ethanol consumption (**Fig. 3K, 4E**) in both sexes, which is in line with the human literature supporting that the anatomy and function of the aIC are altered in AUD patients of both sexes. In contrast to aIC, less information is known about the role of the pIC in AUD patients. Only two studies reported decreased gray matter volume in AUD men and women (56) and elevated resting-state activity in AUD women (55), making it difficult to draw strong conclusions regarding its potentially sexually dimorphic role in AUD. In our study, we identify sex differences in pIC function during persistent ethanol intake in mice, laying the groundwork for future translational investigations of this subregion in men and women suffering from AUD.

## Conclusions

To conclude, our study demonstrated that aIC and pIC play different roles in drinking behaviors. While right aIC glutamatergic neurons encode binge and persistent ethanol drinking independently of sex, right pIC glutamatergic neurons are active during persistent ethanol consumption exclusively in females. In line with these results, inhibition of posterior insular cortex (pIC) excitatory neurons selectively reduced intake of ethanol adulterated with quinine in female mice, indicating a sex-specific role of pIC glutamatergic neurons in mediating persistent ethanol consumption despite aversion. In contrast, inhibition of aIC glutamatergic neurons reduced bitter solution drinking independently of sex. Altogether, our results provide a starting point to characterize further the role of aIC and pIC circuits in sex-dependent addictive-related behaviors.

## ACKNOWLEDGEMENTS

The authors acknowledge the entire Beyeler Lab. We thank Harold Haun and Irene Lorrai for their helpful feedback on setting up the drinking in the dark model. We thank Guillaume Ferreira for the interesting discussions. We thank Sara Laumond, Julie Tessaire, Nathalie Aubailly, and the technical staff of the animal housing facility of the Neurocentre Magendie (INSERM U1215) for their invaluable support.

We acknowledge the support of the Région Nouvelle-Aquitaine, the ‘Avenir program’ of the French Institute of Health (INSERM), the ‘Fondation NRJ-Institut de France’, the ‘Agence Nationale pour la Recherche’ (ANR), the ‘Fondation Schlumberger pour l’Education et la Recherche’ (FSER), and the ‘Fondation Bettencourt-Schueller’ to AB. We are grateful for the support of the ‘Fondation pour la Recherche Médicale’ (FRM) to AB, CN (ARF201909009147) and CF (FDT202404018152), as well as the ‘ATIP program’ (CNRS) to CN. Finally, this research was supported by the French Addictions Fund (Fonds de lutte contre les addictions) under the 2024 call for proposals on Addictive Behaviors and Drugs (CAD): Prevention, Mechanisms, Identification, and Support, jointly led by IReSP and INCa (CAD-V124-014) to CN.

## CONFLICT OF INTEREST

The authors declare that they do not have any competing interests or conflicts of interest (financial or non-financial) related to the material presented in this manuscript.

## Notes

### Competing Interest Statement

The authors have declared no competing interest.

### Summary of Updates

This version has been revised to include new results and updated data analyses.

