## Supplemental file for "Sex differences in insular cortex function in persistent alcohol drinking despite aversion in mice"

### SUPPLEMENTARY INFORMATION

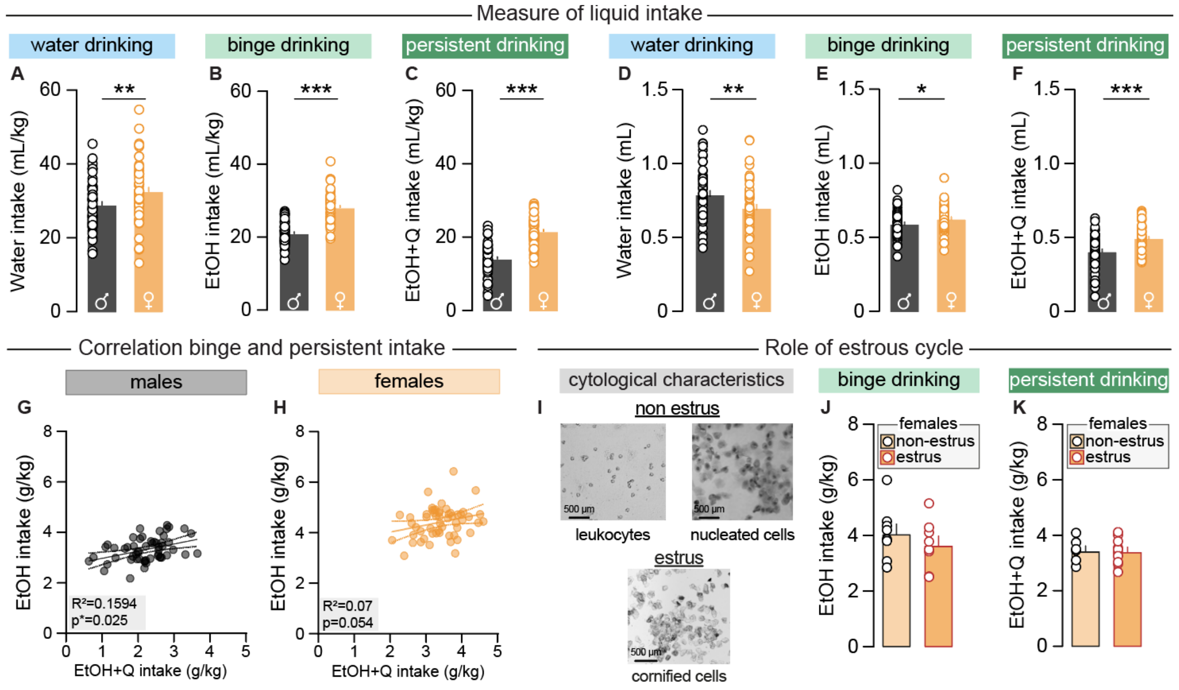


**Fig. S1. Sexual dimorphism of binge and persistent ethanol drinking. A-C** Average of water (A), ethanol (B) and ethanol+quinine (C) intake in mL/kg in male (n=55 for all liquids) and female (water and ethanol n=56, ethanol+quinine n=55) during cycle 1-2 and cycle 3 respectively. **D-F** Average of water (D), ethanol (E) and ethanol+quinine (F) intake in mL in male (n=55 for all liquids) and female (water and ethanol n=56, ethanol+quinine n=55) during cycle 1-2 and cycle 3 respectively. **G, H** correlation between average ethanol consumption during cycle 1-2 and the average ethanol+quinine drinking during cycle 3 in male (G, n=55) and female (H, n=55) mice. **I** Representative images of cells characterizing each phase of the estrous cycle. **J, K** Average of ethanol (J) and ethanol+quinine (K) intake in non-estrus (ethanol n=8, ethanol+quinine n=6) and estrus (ethanol n=8, ethanol+quinine n=10) female mice. Data are shown as mean ± SEM. *p<0.05 **p<0.01 ***p<0.001 represent significant t-test or significant Pearson correlation.

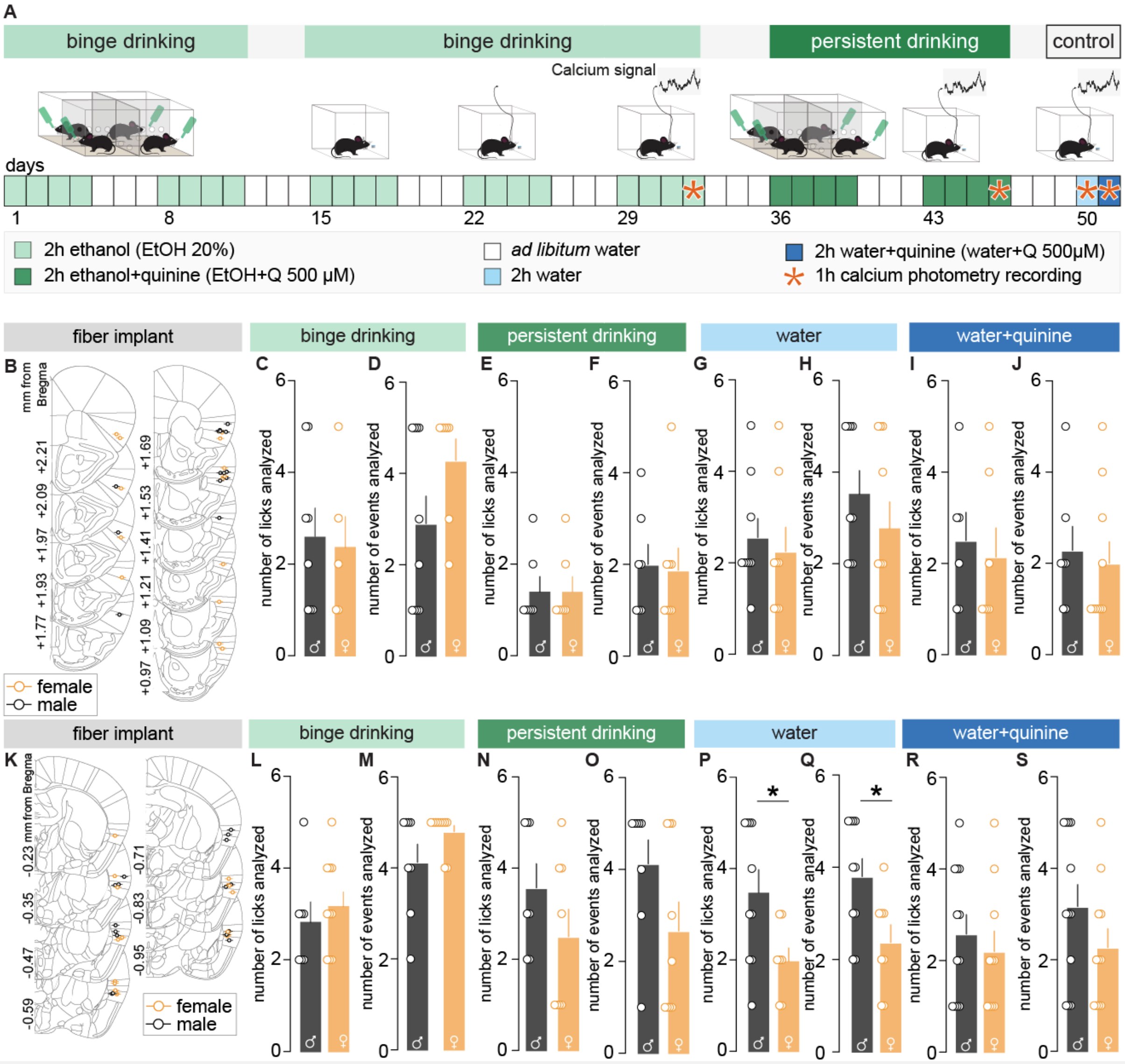


**Fig. S2**. **Experimental procedure of calcium photometry recordings and licking behavior.** **A** Behavioral Timeline. Fiber photometry recordings were performed for 1-hour on the last day of binge (day 32) and persistent (day 46) ethanol drinking cycle, and during a water (day 50) and water+quinine (day 51) single session. **B** Location of fiber implant in the aIC of male and female mice included in the neural activity analysis. **C, D** Average number of single licks (C) and single licks+bouts (D) performed during the 1-hour recording of aIC activity during ethanol licking in males and females. **E, F** Average number of single licks (E) and licks+bouts (F) performed during the 1-hour recording of aIC activity during ethanol+quinine licking in males and females. **G, H** Average number of single licks (G) and licks+bouts (H) performed during the 1-hour recording of aIC activity during water licking in males and females. **I, J** Average number of single licks (I) and licks+bouts (J) performed during the 1-hour recording of aIC activity during water+quinine licking in males and females. **K** Location of fiber implant in the pIC of male and female mice included in the neural activity analysis. **L, M** Average number of single licks (L) and single licks+bouts (M) performed during the 1-hour recording of pIC activity during ethanol licking in males and females. **N, O** Average number of single licks (N) and licks+bouts (O) performed during the 1-hour recording of pIC activity during ethanol+quinine licking in males and females. **P, Q** Average number of single licks (P) and licks+bouts (G) performed during the 1-hour recording of pIC activity during water licking in males and females. **R, S** Average number of single licks (R) and licks+bouts (S) performed during the 1-hour recording of pIC activity during water+quinine licking in males and females. Data are shown as mean ± SEM. *$p<0.05$ significant Student t-test or Mann-Whitney test.

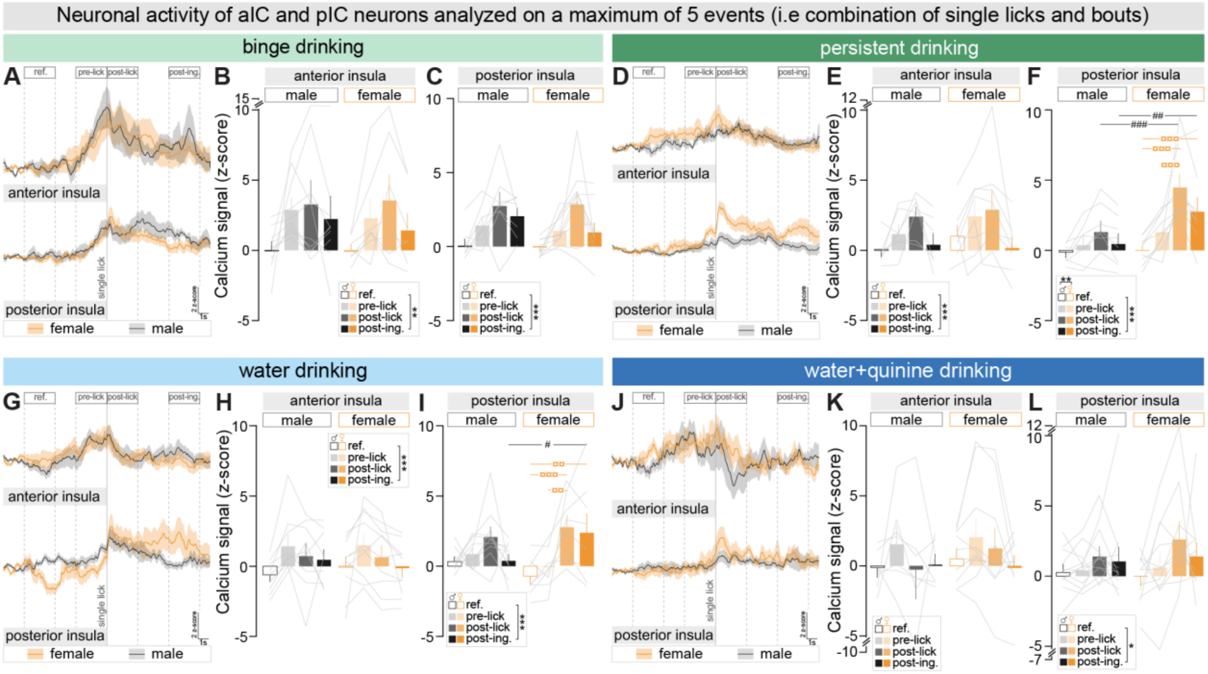


**Fig. S3. Coding properties of anterior (aIC) and posterior (pIC) insular cortex excitatory neurons during binge and persistent ethanol drinking based on a maximum of 5 events. A** Peri-ethanol licking analysis of the calcium signal in the aIC (up) and pIC (bottom) during a maximum of 5 events (licks or bouts) in male (n=10 for aIC and n=8 for pIC) and female mice (n=7 for aIC and n=10 for pIC). **B, C** Average of calcium signal in the aIC (B) or pIC (C) during ethanol licking for the reference, pre-lick, post-lick, and post-ingestive periods in male (n=10 for aIC and n=8 for pIC) and female (n=7 for aIC and n=10 for pIC) mice. **D** Peri-ethanol+quinine licking analysis of the calcium signal in the aIC (up) and pIC (bottom) during a maximum of 5 events in male (n=7 for aIC and n=8 for pIC) and female mice (n=8 for aIC and n=9 for pIC). **E, F** Average of calcium signal in the aIC (E) or pIC (F) during ethanol+quinine licking for the reference, pre-lick, post-lick, and post-ingestive periods in male (n=7 for aIC and n=8 for pIC) and female (n=8 for aIC and n=9 for pIC) mice. **G** Peri-water licking analysis of the calcium signal in the aIC (up) and pIC (bottom) during a maximum of 5 events in male (n=9 for aIC and n=10 for pIC) and female mice (n=10 for aIC and 8 for pIC). **H, I** Average of calcium signal in the aIC (H) or pIC (I) during water licking for the reference, pre-lick, post-lick, and post-ingestive periods in male (n=9 for aIC and n=10 for pIC) and female (n=10 for aIC and n=8 for pIC) mice. **J** Peri-water+quinine licking analysis of the calcium signal in the aIC (up) and pIC (bottom) during a maximum of 5 events (licks or bouts) in male (n=7 for aIC and n=12 for pIC) and female mice (n=10 for aIC and n=11 for pIC). **K, L** Average of calcium signal in the aIC (K) or pIC (L) during water+quinine licking for the reference, pre-lick, post-lick, and post-ingestive periods in male (n=7 for aIC and n=12 for pIC) and female (n=10 for aIC and n=11 for pIC) mice. Data are shown as mean ± SEM. *p<0.05 **p<0.01 ***p<0.001 represent significant main effect of the 2-way RM ANOVA. #p<0.05, ##p<0.01, ###p<0.001, ¤¤p<0.01, ¤¤¤p<0.01 represents a significant Bonferroni post-hoc test.

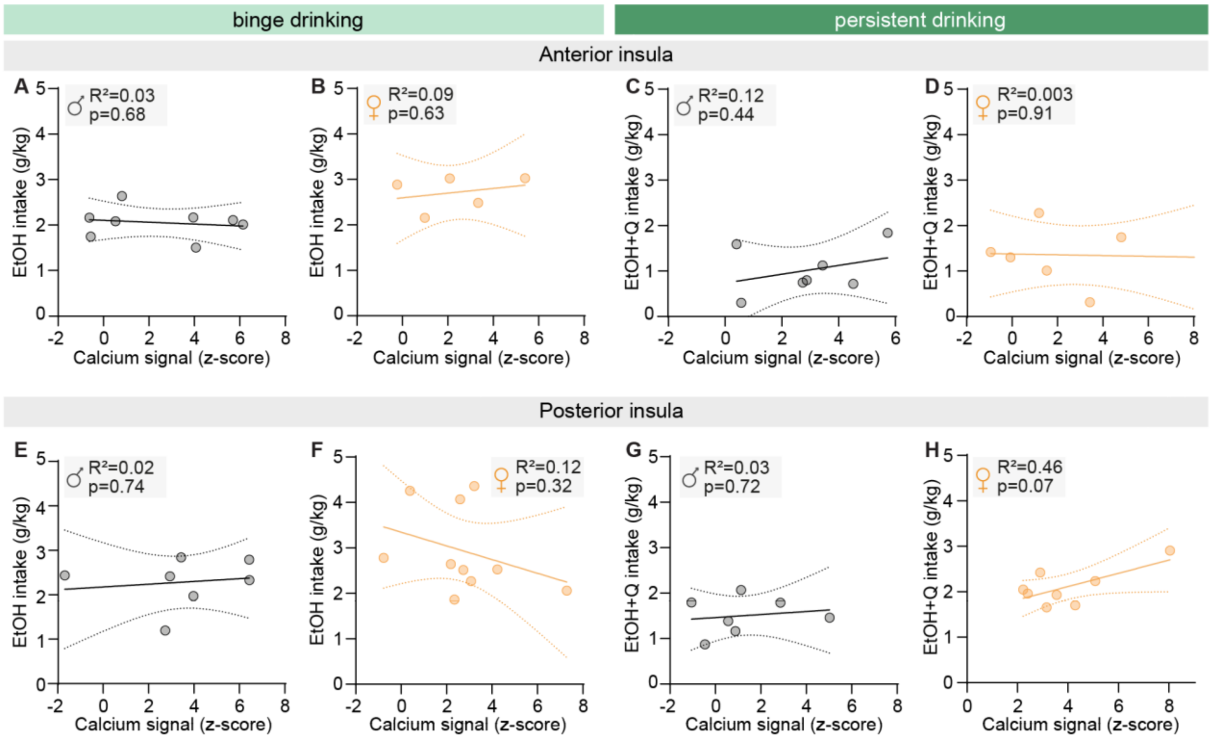


**Fig. S4. Correlation between ethanol and ethanol+quinine intake history and the respective neuronal activity of the anterior (aIC) or posterior (pIC) insular cortex. A, B** Correlation between the average ethanol intake during all the sessions except the recording day and the aIC calcium signal during single licks of ethanol in males (A, n=8) and females (B, n=5). **C, D** Correlation between the average of ethanol+quinine intake during all the sessions except the recording day and the aIC calcium signal single licks of ethanol+quinine in males (C, n =7) and females (D, n=6). **E, F** Correlation between the average ethanol intake during all the sessions except the recording day and the pIC calcium signal during single licks of ethanol in males (E, n=7) and females (F, n=10). **G, H** Correlation between the average of ethanol+quinine intake during all the sessions except the recording day and the pIC calcium signal single licks of ethanol+quinine in males (G, n =7) and females (H, n=8).

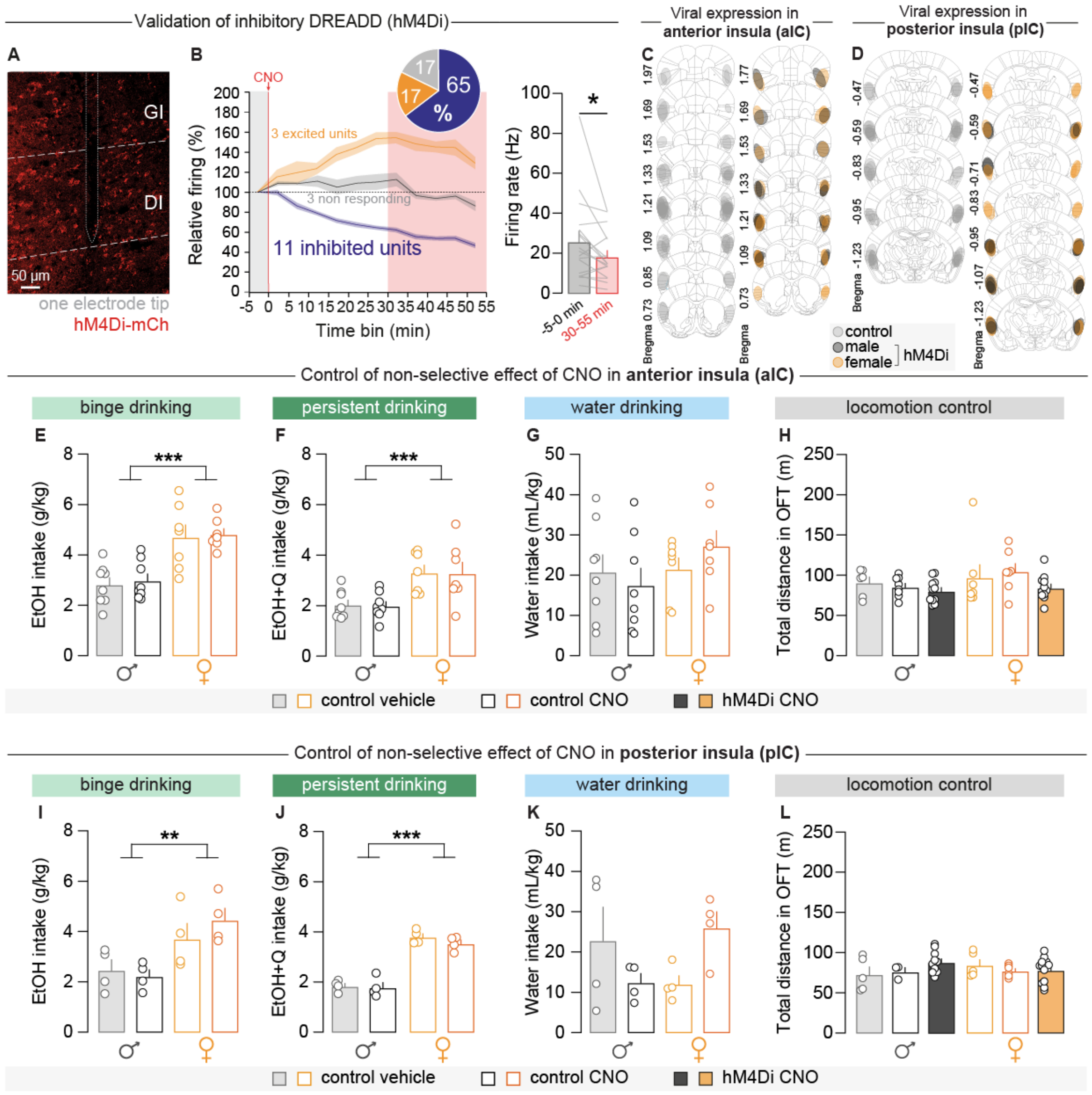


**Fig. S5. Electrophysiological confirmation of neuronal inhibition by hM4Di, histological verification of viral injections and control of clozapine-N-oxyde (CNO) non-specific effects.** **A** Representative image of hM4Di-mCherry expression and a recording track (dashed lines) in the aIC. **B** Percentage of relative firing during chemogenetic inhibition (n=1 mouse, 17 units) for each unit classified as: excited (n=3, 17%), non-responding (n=3, 17%) and inhibited (n=11, 65%). CNO was injected intraperitoneally 5 minutes after the beginning of the recording session (1-h recording session). The 5-min bin before CNO injection corresponds to the baseline. The firing rate decreased during the 30-55 min (red) period when compared to the baseline. **C, D** Location of mCherry and hM4Di-mCherry expression in the aIC (C) or pIC (D) of male and female mice. **E-G** Average of ethanol (E), ethanol+quinine (F) and water (G) intake in aIC control male (n=8) and female (n=7) mice injected with vehicle or CNO. **H** Average of total distance traveled in the open field over 15 minutes in control mice injected with vehicle (n=6 for male, n=7 for female) or CNO (n=8 for male, n=7 for female) and mice expressing the hM4Di receptor injected with CNO (n=11 for male and female). **I-K** Average of ethanol (I), ethanol+quinine (J) and water (K) intake in pIC control male (n=4) and female (n=4) mice injected with vehicle or CNO. **L** Average of total distance traveled in the open field over 15 minutes in control mice injected with vehicle (n=4 for male and female) or CNO (n=4 for male and female) and mice expressing the hM4Di receptor injected with CNO (n=4 for male and female). Data are shown as mean ± SEM. *p<0.05 **p<0.01 ***p<0.001 represent significant t-test and main effect of 2-way ANOVA.

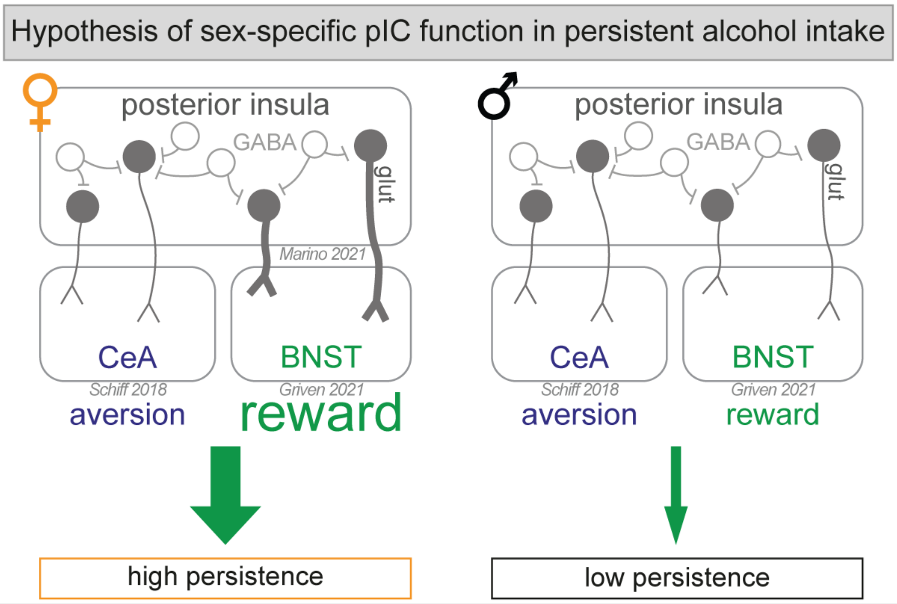


**Fig. S6. Theoretical circuit mechanism underlying sex-specific posterior insula (pIC) function in persistent alcohol drinking based on our data and the current literature. Left panel**: potentiation of the rewarding pIC-BNST glutamatergic pathway by ethanol drinking in female mice, leading to an increased value of the rewarding modality of ethanol over quinine aversion and thus high drinking and persistence toward ethanol. **Right panel**: no potentiation of the rewarding pIC-BNST glutamatergic pathway by ethanol drinking in male mice, leading to balanced reward and aversion pathways, inducing low drinking and low persistence.

### SUPPLEMENTARY METHODS

#### Subjects and housing conditions

The animals used in this study were male (n=85) and female (n=87) C57BL/6J mice aged 10 weeks at the beginning of the experiments (Charles River Laboratory, France). Upon arrival, the mice were housed by sex, 2 to 4 animals per cage, and were provided with *ad libitum* food and water. One week before the beginning of the DID protocol, mice were housed in groups of four separated with transparent perforated dividers allowing them to see and touch each other (**Fig. 1A**). This housing condition allowed to measure individual consumption without isolating the mice which has been shown to increase binge drinking in males and females (1). The animals were housed in the animal facility in a reversed 12-hour light/12-hour dark cycle (8:30 a.m./20:30 p.m. darkness) in controlled conditions of temperature (range 20- 24°C) and hygrometry (range 45-65%). Animal maintenance, treatments and experimental procedures were conducted according to French governmental regulations, and approved by the ethical committee and the Ministry of Education, Research and Innovation (Saisines #22122, #53819) in accordance with the guidelines of the European Communities Council Directives.

#### Stereotaxic surgeries

Animals were anesthetized with isoflurane (Vetflurane®, Virbac, Nice, France, 5% induction and 1.5% maintenance). A subcutaneous injection of Meloxicam (Metacam® 5 mg/kg, Boehringer Ingelheim, Toulouse, France) was performed, eyes were protected by Ocry-gel (Fendigo, Brussels, Belgium) and animals were placed on a heating pad maintained at 39°C during the entire procedure. The upper part of the head was shaved, and an application of local anesthetic (Lidocaine, Zentiva®, Paris, France) was done. The head was fixed on the stereotaxic frame, and disinfected with 10% dermal Betadine and 70% ethanol before incising the skin and periosteum tissue. Craniotomy with stereotactic coordinates in reference to the Bregma was performed at the level of aIC or pIC. The viral vectors were injected using a glass micropipette (3-000-203-G/X, Drummond Scientific Company, Broomall, Pennsylvania, USA) made by a puller (PC-100, Narishige, Setagaya, Japan) and a Nanoliter 2020 injection system

(World Precision Instruments, Sarasota, Florida, USA). All the viral vectors used required 4 weeks to be expressed by the neurons.

Surgeries for *in vivo* extracellular recordings. Mice were unilaterally implanted in the right aIC with a 4x4 fixed array (Innovative Neurophysiology, Bardstown, Kentucky, USA) targeted to the following coordinates: 1.7 mm anterior to Bregma; 3 mm lateral to the midline; and 1.6 mm ventral to the cortical surface. These electrodes consisted in a 16-wire electrode array, arranged in a 4x4 grid (Tungsten wires, 35 µm diameter, 150 µm row spacing, Impedances <1MOhm). The wires were attached to an 18-pin nanoconnector (Omnetics: A79014-001, Minneapolis, Minnesota, USA). Additionally, electrodes were referenced via a silver ground placed above the right visual cortex V1. All implants were secured using Super-Bond cement (Sun Medical, Bois-Guillaume, France). After surgery, mice were allowed to recover for 4 days and were habituated to handling and headstage connection for 3 days. Electrodes were connected to a headstage containing sixteen unity-gain operational amplifiers (Intan: RHD2132, Los Angeles, California, USA). Spiking activity was digitized at 30 kHz and bandpass filtered from 300 Hz to 6 kHz. Before implantation, electrodes were coated with fluorescent tracer (CTB-AF555, Addgene, Watertown, Massachusetts, USA) to allow the tracking of the electrodes.

**Estrous cycle monitoring**

For one session of binge and persistent ethanol intake, we performed vaginal swabs on females to determine the estrous cycle phases (estrus, diestrus, proestrus) based on the cytoarchitecture of the collected cells (**Supplementary Fig. 1I**). Male mice were similarly handled to avoid any bias related to the experimenter's manipulation. Conventionally in the literature, when no addictive-related behaviors differences were observed between diestrus and proestrus these two groups were pooled as non-estrus (2,3), we performed the analysis accordingly.

**In vivo electrophysiological recordings**

Animal. 1 male C57Bl/6J (Charles River, Saint-Germain-Nuelles, France) was used to study the effect of chemogenetic inhibition on the single-unit firing in the aIC. This mouse was previously used in the chemogenetic experiment manipulating aIC activity (**Fig. 4**) and was 16 weeks old at the time of the recordings.

Single-unit recording. Single-unit activity of aIC neurons was recorded in freely moving mice during a 1h session in an open-field arena (60x60cm). Extracellular single-unit recordings signals were acquired with an Open Ephys acquisition board and synchronized with behavior using an electronically controlled camera. Mouse was injected intraperitoneally with clozapine-N-oxide (CNO; 3 mg/kg, Tocris, Bristol, United-Kingdom) 5 minutes after the beginning of the session. The first 5-min bin was used as baseline.

Single-unit analyses. Single-unit action potentials were isolated and analyzed using a custom Python script. Spike detection and sorting were performed with the semi-automatic spike sorter Tridesclous9. A group of action potential waveforms was considered to be generated from a single neuron if the waveforms formed a discrete, isolated cluster in the principal-component space and had a refractory period longer than 1 ms. Unit isolation was verified with auto- and cross-correlation histograms. Units were classified as either excited or inhibited neurons if their firing rate was respectively above or below the average firing in the 5 min baseline + (or -) 2 standard deviations (SD) of this average. Units neither classified as excited nor inhibited have been classified as non-responding neurons. After the recording sessions, the location of the electrodes within the aIC, and the viral expression was controlled using the Paxinos and Franklin atlas.

**Coding properties of aIC and pIC excitatory neurons during binge and persistent ethanol drinking**

The aIC and pIC fiber photometry recordings were performed in different batches. All batches included similar numbers of male and female mice. During fiber photometry recordings, we used polymodal chambers equipped with a drinking sprout controlled by Imetronic interface

and software (Imetronic®, Marcheprime, France). The polymodal chambers (550x450x390mm) are composed of a grid floor, a red house light and a drinking spout connected with a silicon tubing to a 20 mL syringe inserted into a pump. The number of licks was recorded by the Imetronic interface. At the beginning of the recording session, the pump was activated to fill the drinking sprout with 0.114 mL of the appropriate solution. Every time mice performed 10 licks, the pump delivered 0.038 mL to refill the drinking sprout to ensure that it was always full for *ad libitum* access through the session. The mice were handled and connected to the patch cord for 30 minutes in their home cage at least for 3 days before the beginning of the sessions in the multifunctional boxes. During the first binge cycle in the multifunctional boxes, mice were not connected to the patch cord to habituate them to drink in this new environment. For all the following cycles in these boxes, mice were connected to the patch cord for the entire 2-hour sessions, recordings were performed only for 1-hour on recording days 32, 46, 50 and 51 (**Supplementary Fig. S2A**). The investigator was not blinded to the brain region recorded (aIC or pIC) as animals were identified by a unique ID number right after they underwent stereotaxic surgeries, before the beginning of the DID protocol.
<u>Fiber photometry recordings.</u> The optical fiber implanted in pIC transmitted the light emitted by the LEDs (Light-Emitting Diodes) to excite the calcium sensor GCaMP6f. A 20x lens is connected to a cable connected to the optical fiber implanted in the brain and a CMOS (Complementary Metal-Oxide Semiconductor) camera that detects the fluorescence emitted by GCaMP6f, which is indicative of the calcium levels and the neuronal activity *in vivo*. The isosbestic 415 nm channel was used as a negative control, since it is independent of calcium concentration, and allowed to remove motion-related artifacts and signals unrelated to neuronal activity. The power of each channel was set at the maximum power 30 minutes to 1 hour before starting the session to remove patch cord autofluorescence.

**Chemogenetic inhibition of aIC or pIC excitatory neurons during binge and persistent ethanol drinking**
Animals underwent the drinking in the dark (DID) procedure as described in the main manuscript. The aIC and pIC manipulations were performed in different batches. All batches included the similar numbers of male and female mice. To control potential non-selective effects of CNO, mice expressing the control viral vector (AAV9/2-mCaMKII-mCherry-WPRE, ETH Zürich, Switzerland) received CNO (i.p. 3 mg/kg) or vehicle injection 30 minutes before the ethanol, ethanol+quinine and water session. Control mice assigned to vehicle or CNO groups had a similar average of ethanol intake across the sessions (data not shown). Then, as no differences in liquid intake were observed between control mice treated with vehicle or CNO (**Supplementary Fig. S5E-G, S5I-K**), we pooled them as control groups in the main analysis (**Fig. 4**). The investigator was not blinded to the brain region manipulated (aIC or pIC) as animals were identified by a unique ID number right after they underwent stereotaxic surgeries, before the beginning of the DID protocol. Moreover, the investigator needed to know the viral vector injected (hM4Di or control) during stereotaxic surgery to inject the appropriate solution (i.p. vehicle or CNO) at the moment of the experiment.

**Open Field Test**
To control that CNO injection did not impair locomotion, we performed an Open Field Test (OFT, 60x60 cm) at the end of the DID protocol. The OFT was performed during the dark phase of the cycle under red light (15 ± 3 lux at the corners of the OFT), thirty minutes after an injection of CNO (i.p. 3 mg/kg) or vehicle (10 mL/kg) in control and hM4Di animals. Each animal was placed in the same corner of the arena at the beginning of the test and explored for 15 minutes. A camera was placed above the arena to record animal movements.
<u>Data analysis.</u> Bonsai software was used to track the position of the mouse in the arena. A custom-made Python code was used to calculate the distance traveled in the arena during the 15 minutes test.

**Brain extraction, histology and imaging**

At the end of the experimental procedure the animals were anesthetized by intraperitoneal injection of pentobarbital (Exagon® 300 mg/kg, Axience, Pantin, France) and lidocaine (Lidor® 30 mg/kg, Axience, Pantin, France) followed by an intracardiac infusion of Ringer's solution and 4% paraformaldehyde (PFA, antigenfix, F/P0014, MM France, Brignais, France). Brains were extracted, post-fixed in PFA at 4°C overnight and transferred into 30% PBS-sucrose solution for cryoprotection. Then, the brains were embedded in optimum cutting temperature compound (OCT, Cell Path, Newtown, United-Kingdom) and sliced into 100 µm coronal slices, in a freezing sliding microtome (12062999, Thermo Scientific, Bordeaux, France). Sections were incubated with a DNA-specific fluorescent probe Hoechst 33342 (1:5000; Fisher scientific, Bordeaux, France) for 5 minutes, washed 3 times for 10 minutes with PBS 1X and mounted on gelatinized slides. After drying, slides were coverslipped using Polyvinyl alcohol PVA-DABCO mounting medium (Sigma Aldrich, Saint-Louis, Missouri, USA). Images were acquired with fluorescence microscope (THUNDER Imager 3D Tissue, Leica, Wetzlar, Germany) with a 10X objective and a Retiga 2000R CCD camera (QImaging, Surrey, Canada) to control the viral vector expression level, the localization of the viral expression and the optic fiber implantation. Mice were excluded from the analysis when optic fiber placement or viral expression patterns were out of the brain target.

**Statistical analysis**

All statistical analyses were done using GraphPad Prism 9 software (GraphPad Software Inc.). The values are represented according to the mean ± SEM. The kinetics of liquid intake were compared statistically using 2-way RM-ANOVA (factors: sex and sessions). After performing a normality Gaussian test (Shapiro-Wilk test), the comparison of the liquid intake average during the sessions was performed using an unpaired parametric two-tailed t-test if the data distribution followed a normal distribution, otherwise an unpaired (Mann-Whitney) or paired (Wilcoxon) non-parametric two-tailed t-test was used. For the analysis of the average liquid intake, blood ethanol concentrations, and licking bouts during photometry recordings (**Fig. 1B, C, E, F, Supplementary Fig. S1A-F, J, K, Supplementary Fig. S2C-J, S2L-S**) we used a two-tailed t-test. For liquid intake during chemogenetic tests, we used a 2-way RM-ANOVA (factors: sex, virus). For fiber photometry analysis, the calcium signal changes between the reference and licking period in males and females was analyzed using a 2-way ANOVA (factors: sex, period) for all liquids. For 2-way ANOVAs, we followed up on significant interaction effects with a Bonferroni post-hoc test except for **Fig. 4E, G** where the interaction was approaching significance therefore we performed an exploratory Bonferroni post-hoc analysis for the main effect. Detailed statistical analyses are described in the Results section. Our multifactorial ANOVAs yielded multiple main and interaction effects. Thus, we only reported significant effects that are critical for data interpretation (see Table S1 for a complete statistical reporting). The sample sizes were chosen to match those reported in previous publications and the appropriate statistical comparisons were performed.

| Figure No | Test used | Factor names | Values | p-Value |
|---|---|---|---|---|
| Figure 1B | Unpaired Student t-test | Sex | t=3.864, df=24 | 0.0007 |
| Figure 1C | Unpaired Student t-test<br>Fischer's exact test | Sex<br>Sex | t=0.2454, df=24<br>males 3/10<br>females 2/11 | 0.8082<br><br>>0.9999 |
| Figure 1D | 2-way ANOVA | Sex<br>ethanol proportion | $F_{1, 60}$=0.000<br>$F_{1, 60}$=132.2 | p>0.999<br>p<0.0001 |
| Figure 1E | Unpaired Student t-test | Sex | t=3.679, df=30 | 0.0009 |
| Figure 1F | Unpaired Mann-Whitney test<br>Fischer's exact test | Sex<br>Sex | U=71<br>males 6/10<br>females 12/4 | 0.0318<br><br>0.0732 |
| Figure 2B | 2-way RM-ANOVA | Sex<br>Session<br>Sex x session | $F_{1, 109}$=7.208<br>$F_{3, 327}$=3.342<br>$F_{3, 327}$=0.1697 | 0.0084<br>0.0195<br>0.9168 |
| Figure 2C | Unpaired Student t-test | Sex | t=2.685, df=109 | 0.0084 |
| Figure 2D | 2-way RM-ANOVA | Sex<br>Session<br>Sex x session | $F_{1,109}$=116.7<br>$F_{5.331, 581.1}$=3.577<br>$F_{6, 654}$=0.9494 | <0.0001<br>0.0027<br>0.4589 |
| Figure 2E | Unpaired Student t-test | Sex | t=10.80, df=109 | <0.0001 |
| Figure 2F | 2-way RM-ANOVA | Sex<br>Session<br>Sex x session | $F_{1,108}$=103.4<br>$F_{1.887,203.8}$=4.341<br>$F_{2, 216}$=1.951 | <0.0001<br>0.0159<br>0.1446 |
| Figure 2G | Unpaired Student t-test | Sex | t=10.17, df=108 | <0.0001 |
| Figure 2H | 2-way RM-ANOVA | Sex<br>Session<br>Sex x session | $F_{1,46}$=20.64<br>$F_{6. 276}$=23.41<br>$F_{6, 216}$=3.450 | <0.0001<br><0.0001<br>0.0027 |
| Figure 2I | 2-way RM-ANOVA | Sex<br>Liquid<br>Sex x liquid | $F_{1, 46}$=17.85<br>$F_{1, 46}$=85.86<br>$F_{1, 46}$=9.787 | 0.0001<br><0.0001<br>0.0030 |
| Figure 3E | Unpaired Mann-Whitney test | Sex | U=145 | 0.0143 |
| Figure 3F | Unpaired Student t-test | Sex | t=2.357, df=41 | 0.0233 |
| Figure 3H | 2-way RM-ANOVA | Sex<br>Period<br>Sex x period | $F_{1, 11}$=0.2616<br>$F_{3, 33}$=7.374<br>$F_{3, 33}$=0.3421 | 0.6191<br>0.0007<br>0.7950 |
| Figure 3I | 2-way RM-ANOVA | Sex<br>Period<br>Sex x period | $F_{1, 15}$=2.916<br>$F_{3, 45}$=9.974<br>$F_{3, 45}$=2.875 | 0.1083<br><0.0001<br>0.0465 |
| Figure 3K | 2-way RM-ANOVA | Sex | $F_{1, 12}$=0.1030 | 0.7538 |

| | | | | |
|---|---|---|---|---|
| | | Period<br>Sex x period | $F_{3, 36}$=4.757<br>$F_{3, 36}$=0.7980 | 0.0068<br>0.5031 |
| Figure 3L | 2-way RM-ANOVA | Sex<br>Period<br>Sex x period | $F_{1, 13}$=3.447<br>$F_{3, 39}$=11.75<br>$F_{3, 39}$=3.717 | 0.0862<br><0.0001<br>0.0192 |
| Figure 3N | 2-way RM-ANOVA | Sex<br>Period<br>Sex x period | $F_{1, 15}$=0.09366<br>$F_{3, 45}$=3.956<br>$F_{3, 45}$=0.1703 | 0.7683<br>0.0138<br>0.9159 |
| Figure 3O | 2-way RM-ANOVA | Sex<br>Period<br>Sex x period | $F_{1, 15}$=1.101<br>$F_{3, 45}$=6.288<br>$F_{3, 45}$=2.814 | 0.3107<br>0.0012<br>0.0498 |
| Figure 3Q | 2-way RM-ANOVA | Sex<br>Period<br>Sex x period | $F_{1, 11}$=0.9428<br>$F_{3, 33}$=2.186<br>$F_{3, 33}$=0.1650 | 0.3524<br>0.1082<br>0.9192 |
| Figure 3R | 2-way RM-ANOVA | Sex<br>Period<br>Sex x period | $F_{1, 20}$=0.9419<br>$F_{3, 60}$=2.962<br>$F_{3, 60}$=0.8110 | 0.3434<br>0.0392<br>0.4928 |
| Figure 4D | 2-way ANOVA | Sex<br>Virus<br>Sex x virus | $F_{1, 49}$=39.66<br>$F_{1, 49}$=0.9415<br>$F_{1, 49}$=2.582 | <0.0001<br>0.3367<br>0.1145 |
| Figure 4E | 2-way ANOVA | Sex<br>Virus<br>Sex x virus | $F_{1, 48}$=15.78<br>$F_{1, 48}$=6.180<br>$F_{1, 48}$=3.151 | 0.0002<br>0.0165<br>0.0822 |
| Figure 4F | 2-way ANOVA | Sex<br>Virus<br>Sex x virus | $F_{1, 48}$=2.416<br>$F_{1, 48}$=0.01232<br>$F_{1, 48}$=0.001176 | 0.1267<br>0.9121<br>0.9728 |
| Figure 4G | 2-way ANOVA | Sex<br>Virus<br>Sex x virus | $F_{1, 25}$=1.541<br>$F_{1, 25}$=14.72<br>$F_{1, 25}$=3.839 | 0.2259<br>0.0008<br>0.0613 |
| Figure 4H | 2-way ANOVA | Sex<br>Virus<br>Sex x virus | $F_{1, 31}$=35.05<br>$F_{1, 31}$=0.2444<br>$F_{1, 31}$=0.06311 | <0.0001<br>0.6245<br>0.8033 |
| Figure 4I | 2-way ANOVA | Sex<br>Virus<br>Sex x virus | $F_{1, 31}$=58.57<br>$F_{1, 31}$=14.24<br>$F_{1, 31}$=5.263 | <0.0001<br>0.0007<br>0.0287 |
| Figure 4J | 2-way ANOVA | Sex<br>Virus<br>Sex x virus | $F_{1, 31}$=1.111<br>$F_{1, 31}$=0.03462<br>$F_{1, 31}$=0.4508 | 0.3000<br>0.8536<br>0.5069 |
| Figure 4K | 2-way ANOVA | Sex<br>Virus<br>Sex x virus | $F_{1, 24}$=6.284<br>$F_{1, 24}$=7.063<br>$F_{1, 24}$=4.582 | 0.0194<br>0.0138<br>0.0427 |
| Figure S1A | Unpaired Student t-test | Sex | t=2.685, df=109 | 0.0084 |

| | | | | |
|---|---|---|---|---|
| Figure S1B | Unpaired Student t-test | Sex | t=10.80, df=109 | <0.0001 |
| Figure S1C | Unpaired Student t-test | Sex | t=10.17, df=108 | <0.0001 |
| Figure S1D | Unpaired Student t-test | Sex | t=2.669, df=109 | 0.0088 |
| Figure S1E | Unpaired Student t-test | Sex | t=2.127, df=109 | 0.0357 |
| Figure S1F | Unpaired Student t-test | Sex | t=4.582, df=108 | <0.0001 |
| Figure S1G | Pearson correlation | Male | $R^2$=0.1594 | 0.0025 |
| Figure S1H | Pearson correlation | Female | $R^2$=0.07033 | 0.0504 |
| Figure S1J | Unpaired Student t-test | Estrous phase | t=0.8986, df=14 | 0.3841 |
| Figure S1K | Unpaired Student t-test | Estrous phase | t=0.1341, df=14 | 0.8952 |
| Figure S2C | Unpaired Student t-test | Sex | t=0.2348, df=11 | 0.8187 |
| Figure S2D | Unpaired Mann-Whitney test | Sex | U=20 | 0.1266 |
| Figure S2E | Unpaired Mann-Whitney test | Sex | U=24.50 | >0.9999 |
| Figure S2F | Unpaired Mann-Whitney test | Sex | U=25 | 0.7737 |
| Figure S2G | Unpaired Mann-Whitney test | Sex | U=28 | 0.4953 |
| Figure S2H | Unpaired Mann-Whitney test | Sex | U=30.50 | 0.2330 |
| Figure S2I | Unpaired Mann-Whitney test | Sex | U=17 | 0.5304 |
| Figure S2J | Unpaired Mann-Whitney test | Sex | U=27.5 | 0.4990 |
| Figure S2L | Unpaired Mann-Whitney test | Sex | U=27.5 | 0.5225 |
| Figure S2M | Unpaired Mann-Whitney test | Sex | U=26 | 0.1833 |
| Figure S2N | Unpaired Mann-Whitney test | Sex | U=16.5 | 0.1520 |
| Figure S2O | Unpaired Mann-Whitney test | Sex | U=21 | 0.1419 |
| Figure S2P | Unpaired Student t-test | Sex | t=2.382, df=15 | 0.0309 |
| Figure S2Q | Unpaired Mann-Whitney test | Sex | U=16 | 0.0300 |
| Figure S2R | Unpaired Mann-Whitney test | Sex | U=51 | 0.5712 |
| Figure S2S | Unpaired Mann-Whitney test | Sex | U=45 | 0.2005 |
| Figure S3B | 2-way RM-ANOVA | Sex<br>Period<br>Sex x period | $F_{1, 15}$=0.05858<br>$F_{3, 45}$=5.067<br>$F_{3, 45}$=0.1410 | 0.8120<br>0.0042<br>0.9349 |
| Figure S3C | 2-way RM-ANOVA | Sex<br>Period<br>Sex x period | $F_{1, 16}$=0.6862<br>$F_{3, 48}$=13.06<br>$F_{3, 48}$=0.5825 | 0.4196<br><0.0001<br>0.6294 |
| Figure S3E | 2-way RM-ANOVA | Sex<br>Period | $F_{1, 13}$=0.5842<br>$F_{3, 39}$=8.703 | 0.4583<br>0.0002 |

| | | | | |
|---|---|---|---|---|
| | | Sex x period | $F_{3, 39}$=0.8453 | 0.4775 |
| Figure S3F | 2-way RM-ANOVA | Sex<br>Period<br>Sex x period | $F_{1, 15}$=6.391<br>$F_{3, 45}$=17.43<br>$F_{3, 45}$=4.158 | 0.0232<br><0.0001<br>0.0110 |
| Figure S3H | 2-way RM-ANOVA | Sex<br>Period<br>Sex x period | $F_{1, 17}$=0.0004177<br>$F_{3, 51}$=6.655<br>$F_{3, 51}$=1.031 | 0.9839<br>0.0007<br>0.3869 |
| Figure S3I | 2-way RM-ANOVA | Sex<br>Period<br>Sex x period | $F_{1, 16}$=0.01279<br>$F_{3, 48}$=9.465<br>$F_{3, 48}$=4.006 | 0.9114<br><0.0001<br>0.0127 |
| Figure S3K | 2-way RM-ANOVA | Sex<br>Period<br>Sex x period | $F_{1, 15}$=0.4734<br>$F_{3, 45}$=2.077<br>$F_{3, 45}$=0.5531 | 0.5019<br>0.1166<br>0.6487 |
| Figure S3L | 2-way RM-ANOVA | Sex<br>Period<br>Sex x period | $F_{1, 21}$=0.1895<br>$F_{3, 63}$=3.576<br>$F_{3, 63}$=0.6354 | 0.6678<br>0.0186<br>0.5950 |
| Figure S4A | Pearson correlation | Male | $R^2$=0.02940 | 0.6847 |
| Figure S4B | Pearson correlation | Female | $R^2$=0.08590 | 0.6323 |
| Figure S4C | Pearson correlation | Male | $R^2$=0.1218 | 0.4430 |
| Figure S4D | Pearson correlation | Female | $R^2$=0.002931 | 0.9082 |
| Figure S4E | Pearson correlation | Male | $R^2$=0.02320 | 0.7444 |
| Figure S4F | Pearson correlation | Female | $R^2$=0.1203 | 0.3262 |
| Figure S4G | Pearson correlation | Male | $R^2$=0.02774 | 0.7211 |
| Figure S4H | Pearson correlation | Female | $R^2$=0.4587 | 0.0650 |
| Figure S5B | Paired Wilcoxon test | Period | | 0.0232 |
| Figure S5E | 2-way ANOVA | Sex<br>Treatment<br>Sex x treatment | $F_{1, 26}$=32.61<br>$F_{1, 26}$=0.1768<br>$F_{1, 26}$=0.005399 | <0.0001<br>0.6776<br>0.9420 |
| Figure S5F | 2-way ANOVA | Sex<br>Treatment<br>Sex x treatment | $F_{1, 26}$=19.22<br>$F_{1, 26}$=0.01932<br>$F_{1, 26}$=0.0002715 | 0.0002<br>0.8905<br>0.9870 |
| Figure S5G | 2-way ANOVA | Sex<br>Treatment<br>Sex x treatment | $F_{1, 26}$=1.785<br>$F_{1, 26}$=0.09212<br>$F_{1, 26}$=1.330 | 0.1931<br>0.7639<br>0.2593 |
| Figure S5H | 2-way ANOVA | Sex<br>Treatment<br>Sex x treatment | $F_{1, 38}$=2.006<br>$F_{2, 38}$=1.043<br>$F_{2, 38}$=0.5823 | 0.1648<br>0.3622<br>0.5635 |
| Figure S5I | 2-way ANOVA | Sex<br>Treatment | $F_{1, 12}$=14.01<br>$F_{1, 12}$=0.2930 | 0.0028<br>0.5982 |

| | | | | |
|---|---|---|---|---|
| | | Sex x treatment | $F_{1, 12}$=1.161 | 0.3024 |
| Figure S5J | 2-way ANOVA | Sex<br>Treatment<br>Sex x treatment | $F_{1, 12}$=150.1<br>$F_{1, 12}$=1.087<br>$F_{1, 12}$=0.4430 | <0.0001<br>0.3176<br>0.5183 |
| Figure S5K | 2-way ANOVA | Sex<br>Treatment<br>Sex x treatment | $F_{1, 12}$=0.07950<br>$F_{1, 12}$=0.1326<br>$F_{1, 12}$=6.324 | 0.7828<br>0.7221<br>0.0272 |
| Figure S5L | 2-way ANOVA | Sex<br>Treatment<br>Sex x treatment | $F_{1, 30}$=0.01501<br>$F_{2, 30}$=0.5902<br>$F_{2, 30}$=1.732 | 0.9033<br>0.5605<br>0.1941 |

**Table S1.** Statistical analysis

| | Anterior insula | | Posterior insula | | | | | |
|---|---|---|---|---|---|---|---|---|
| Experiment | #1 | #2 | #1 | #2 | #3 | #4 | #5 | #6 |
| male | 7 | 7 | 3 | 3 | 3 | 2 | 2 | 3 |
| female | 7 | 7 | 2 | 3 | 2 | 3 | 4 | 3 |

**Table S2:** Number of animals per experimental replication for photometry recording of anterior and posterior insula neuronal activity in male and female mice.

| | Anterior insula | | | | Posterior insula | | | |
|---|---|---|---|---|---|---|---|---|
| | Experiment #1 | | Experiment #2 | | Experiment #1 | | Experiment #2 | |
| | Control (mCherry) | hM4Di | Control (mCherry) | hM4Di | Control (mCherry) | hM4Di | Control (mCherry) | hM4Di |
| male | 10 | 6 | 6 | 5 | 7 | 8 | 6 | 6 |
| female | 7 | 6 | 7 | 6 | 7 | 8 | 7 | 8 |

**Table S3:** Number of animals per experimental replication for chemogenetic inhibition of anterior and posterior insula neurons in male and female mice.

| | Anterior insula | | | | | | | |
|---|---|---|---|---|---|---|---|---|
| | Single licks | | | | First 5 events | | | |
| | Ethanol | Ethanol+Q | Water | Water+Q | Ethanol | Ethanol+Q | Water | Water+Q |
| male | 8 | 7 | 9 | 6 | 10 | 7 | 9 | 7 |
| female | 5 | 7 | 8 | 12 | 7 | 8 | 10 | 10 |

**Table S4:** Number of animals for photometry recording of anterior insula neurons for each analysis (single licks vs events) and each liquid tested.

| | Posterior insula | | | | | | | |
|---|---|---|---|---|---|---|---|---|
| | Single licks | | | | First 5 events | | | |
| | Ethanol | Ethanol+Q | Water | Water+Q | Ethanol | Ethanol+Q | Water | Water+Q |
| male | 7 | 7 | 10 | 7 | 8 | 8 | 10 | 12 |
| female | 10 | 8 | 7 | 10 | 10 | 9 | 8 | 11 |

**Table S5** Number of animals for photometry recording of posterior insula neurons for each analysis (single licks vs events) and each liquid tested.
